# Sensorimotor adaptation and cue reweighting compensate for distorted 3D shape information, accounting for paradoxical perception-action dissociations

**DOI:** 10.1101/540187

**Authors:** Evan Cesanek, Jordan A. Taylor, Fulvio Domini

## Abstract

Reach-to-grasp movements can show surprising accuracy even when the perceived 3D shape of the target is distorted. One explanation of this paradox is that a specialized “vision-for-action” system provides accurate shape estimates by relying selectively on stereo information and ignoring less reliable depth cues like texture and shading. However, the key support for this hypothesis has come from studies that analyze average behavior across many visuomotor interactions where available sensory feedback reinforces stereo information. The present study, which carefully accounts for the effects of feedback, shows that these interactions are actually planned using the same cue-combination function as perception, and that apparent dissociations can arise due to two distinct supervised learning processes: sensorimotor adaptation and cue reweighting. In four experiments, we show that (1) when a distorted depth cue biases perception (*e.g.*, surfaces appear flattened by a fixed amount), sensorimotor adaptation rapidly adjusts motor outputs to compensate for this constant error. However, (2) when the distorted depth cue is unreliable, leading to variable errors across a set of objects (*i.e.*, some depths are overestimated, others underestimated), then the relative weighting of depth cues is gradually adjusted to reduce the misleading effect of the unreliable cue, consistent with previous studies of cue reweighting. In our third experiment, we show that (3) cue reweighting involves correlated changes in perceptual judgments and motor behavior, indicating a close functional link between perception and action, and that (4) cue reweighting occurs only in response to an unreliable depth cue, and not in response to a biased cue, presumably because movement errors caused by biased cues are rapidly resolved by sensorimotor adaptation. Taken together, these findings emphasize the mutual dependency of perception and action, with perception directly guiding action, and actions producing error signals that drive motor and perceptual learning.

**Author summary:** When interacting with 3D surfaces and objects, sensory feedback is available that could improve future performance via supervised learning processes. In four experiments, we confirm that natural visuomotor interactions with 3D objects lead to sensorimotor adaptation and cue reweighting, two distinct learning processes that are uniquely suited to resolve errors caused by biased and noisy processing of visual cues to depth. Whereas sensorimotor adaptation appears to shift motor outputs without altering 3D shape perception, we show that cue reweighting has correlated effects on perceptual judgments and motor behavior. The speed and flexibility of these two forms of learning provides a simple alternative explanation of why perception and action are sometimes found to be dissociated in experiments where some depth cues are consistent with sensory feedback while others are faulty. Rather than having a specialized “vision-for-action” system that relies preferentially on stereo information, as suggested in some current theories, the brain appears to combine multiple depth cues in the same way for perceptual judgments and visuomotor interactions. We propose that the sensory-prediction errors obtained during these interactions, which are widely recognized to drive sensorimotor adaptation, may also be responsible for producing the more gradual perceptual shifts associated with cue reweighting.

## Introduction

Picking up a nearby object is a basic human behavior that involves a surprising level of computational complexity. To shape and orient the hand for a stable grasp, your visual system must first process a diverse assortment of depth cues, then combine these signals into a single estimate of the target object’s 3D shape, which is then transformed into appropriate motor commands. Since the availability and quality of individual depth cues can vary widely from one situation to the next, leading to bias and noise in 3D shape perception, the process of cue combination is one of the most challenging aspects of this problem. Indeed, even when viewing real, fully illuminated objects from within reaching distance, human visual perception often fails to provide veridical estimates of 3D shape [1–3]. This raises the question of how we manage to produce consistently accurate movements despite the variable distortions that afflict 3D shape perception.

A popular explanation of this paradox is that perception and action are supported by separate visual processing of 3D shape information, with the purported “vision-for-action” system capable of recovering more accurate spatial estimates than the “vision-for-perception” system [4]. One cornerstone of support for this theory is the literature regarding the effects of visual illusions on motor behavior, where many studies have reported that motor responses are more accurate than perceptual judgments of the same illusory stimuli. For example, Bruggeman *et al*. [5] presented participants with the Ames window illusion, which is created by putting carefully constructed texture cues specifying a 3D scene in conflict with stereo cues specifying the actual scene, a flat surface. When participants were asked to make perceptual judgments regarding the degree of surface slant, they were misled by the biased texture cues. However, when asked to interact with the Ames window by making bimanual pointing movements targeting its left and right edges, the movements were, on average, more accurate with respect to the physical slant specified by the stereo cue. These findings were interpreted as evidence that motor planning relies preferentially on stereo information.

The Ames window is an example of an experimental stimulus that involves a biased depth cue: illusory texture information consistently indicates that the display is more slanted than it actually is. Meanwhile, other experiments have examined visuomotor responses in situations involving an unreliable depth cue, *i.e.*, one that suddenly becomes less correlated with the physical layout of the environment. For example, Knill [6] rendered slanted surfaces using conflicting stereo and texture information and asked participants to place an object so that its bottom would be parallel with the surface at contact. On some trials, one of the cues was perturbed to specify either more or less slant than the underlying physical surface. So, unlike the study of Bruggeman *et al*. [5], which involved a constant bias in the texture cue (and thus a relatively constant perceptual bias), the errors experienced in Knill’s study [6] were variable, changing sign randomly throughout the task. Yet the results were similar: the average orientation of the handheld object when it made contact with the surface was slightly closer to the stereo slant.

Both of the cited studies, and others like them, have been interpreted as evidence that “vision-for-action” selectively relies on stereo information, enabling the motor system to avoid making mistakes based on faulty processing of other, typically less reliable cues [7]. Stereo information is special, it is argued, because binocular disparities are a straightforward function of object depth, viewing distance, and interpupillary distance. Since interpupillary distance is relatively fixed in adults, an estimate of viewing distance from ocular convergence should be all the visual system needs to recover a metric estimate of object depth. Other cues, like texture, require the visual system to make additional, potentially complex assumptions (*e.g.*, about the process that generated the texture pattern) before it is possible to arrive at any specific metric estimate.

This interpretation originates from the two visual streams theory, which has been broadly influential in perception and action research, explaining a variety of neuropsychological and psychophysical findings [4, 7, 8]. Here, we focus specifically on testing an alternative explanation of the apparent preference for stereo information in visuomotor tasks involving biased and/or unreliable depth cues. Our account eliminates the need to posit separate 3D shape estimates for perception and action, showing instead how some dissociations can be explained as artifacts produced by averaging over many trials where informative sensory feedback is available. We focus on the effects of two supervised learning processes—*sensorimotor adaptation* [9–11] and *cue reweighting* [12–15]—that could shift motor responses in a way that appears to privilege stereo information (or potentially any other cue, depending on the feedback conditions). By accounting for these processes, we are able to show how averaged movement kinematics can appear to combine depth cues differently than perceptual judgments without positing separate cue-combination functions for perception and action.

### Study overview

In Experiment 1, we show that a standard computational model of sensorimotor adaptation can account for changes in the motor response during exposure to a constant bias in stereo information; this is the converse of Bruggeman *et al*.’s experimental design [5], which involved a constant bias in texture information. In Experiment 2, we examine how the motor response changes over time in the more complex scenario where one cue becomes uncorrelated with sensory feedback, producing variable errors. This is similar to Knill’s study [6] and previous studies on cue reweighting in perception [12–14]. Our results show that these two learning processes operate as expected in these situations, supporting our alternative explanation of why goal-directed actions appear to prefer stereo information in studies that provide stereo-consistent feedback.

Additionally, the present experiments extend current knowledge of cue reweighting, a learning process that has been studied considerably less than sensorimotor adaptation. Whereas previous studies on cue reweighting in perception provided haptic information via tightly constrained exploratory hand movements and emphasized explicit intermodal comparisons of vision and touch, Experiment 2 confirms that cue reweighting of motor responses also occurs in natural visuomotor tasks, replicating and extending our findings from a similar experiment [15]. However, since we did not include perceptual tests, Experiment 2 leaves open the possibility that the observed cue reweighting was restricted to the visuomotor system, and thus qualitatively different than the perceptual cue reweighting shown previously [12–14]. This led us to conduct Experiment 3, in which we demonstrate that cue reweighting in the motor response is indeed the result of upstream perceptual changes. Experiment 3 also provided a chance to generalize our findings on cue reweighting to another 3D shape property, object depth. Having already established perceptual [12] and visuomotor cue reweighting of stereo and texture information for slanted surfaces, in Experiment 3 we adopted a new set of stimuli, this time testing for correlated perceptual and motoric changes as participants grasped small 3D objects along their depth dimension. In sum, the findings from this series of experiments reveal the mutual dependency between perception and action, in contrast to the dissociated view of these functions, and demonstrate the multiple ways that feedback from visuomotor interactions is exploited to improve future performance.

## Results

### Experiment 1: Sensorimotor adaptation compensates for a biased depth cue

To motivate our model of sensorimotor adaptation to a biased depth cue, it will be helpful to walk through a short example of motor interaction with a cue-conflict stimulus akin to the Ames window. Consider the stereogram in Fig 1a, which was constructed so that the perceived surface should appear to have a deeper slant than the flat plane of the document it is printed on (*i.e.*, when held at about arm’s length, the top edge should appear slightly beyond the page while the bottom edge is slightly protruding). As depicted in Fig 1b, this perceived slant (yellow) is due to perceptual combination of a stereo slant (red) and a conflicting texture slant (blue) printed on the flat surface of the page (transparent frame). Now, imagine trying to simultaneously place your index finger on the top edge and your thumb on the bottom edge of this surface. If the perceived slant guides visuomotor planning, you will reach with your grip angled slightly forward, so the index finger leads the thumb. However, this means you will bump the physical page sooner than expected with your index finger (Fig 1c), giving rise to an error signal. In the process of sensorimotor adaptation, this error signal is exploited to adjust the mapping from visually perceived slants to motor outputs so that the next movement will be more appropriate for the physical slant of the surface. As you repeatedly interact with this surface, error corrections accumulate, so after a few trials all of your reaches will tend to be accurate.

**Fig 1.**
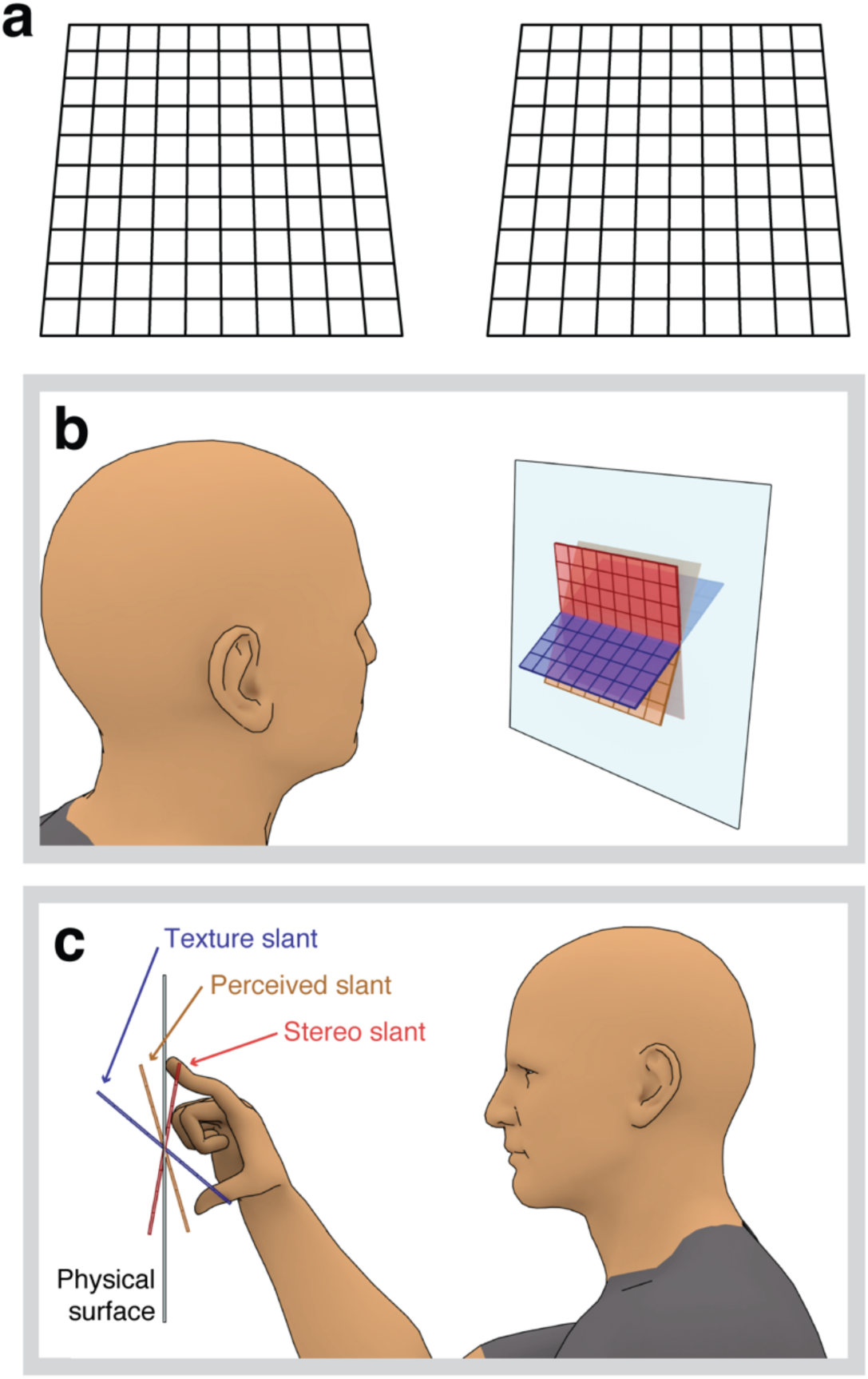
Interacting with slanted surfaces defined by conflicting stereo and texture cues. (a) Example stimulus (cross-fuse). Stereo information specifies a surface with the top edge nearer to the viewer than the bottom edge, while texture information specifies an opposite slant direction. Observers typically perceive a slant that is between the two component slants. (b) An observer viewing the cue-conflict slant stimulus rendered in panel a. The physical surface of the page is depicted by the large, transparent frame. The component stereo and texture surfaces are shown in conflict, with the perceived slant in the middle. In this example of viewing a stereogram printed on a flat page, the physical surface does not match the perceived slant, stereo slant, or the texture slant. (c) In our experiments, the observer attempts to place the index finger and thumb simultaneously on the displayed surface, as shown. When the planned grip orientation is not appropriate for the physical slant, an error signal is registered as one of the fingers bumps into the surface earlier than anticipated. We hypothesize that during this Grip Placement task, the grip orientation will initially target the perceived slant, but gradually come to target the haptically reinforced slant of the underlying physical surface. Unlike the example depicted here, the physical surfaces in our experiments were made to be consistent with either texture or stereo information, depending on the condition.

Sensorimotor adaptation to a constant bias in slant perception can be formalized using a linear state-space model of proportional error correction [16, 17]. On a given trial *n*, the observer perceives some slant *ŝ* that is a function of the available texture and stereo information *Ψ*(*s_T_, s_S_*):

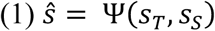

The planned grip orientation *y_n_* for that trial is the combination of the perceived slant *ŝ* and an adjustable internal state *x_n_*:

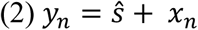

When the planned grip orientation *y_n_* does not match the physical surface slant *s_Φ_*, haptic feedback produces an error signal *ε_n_* as a function of the difference between the planned grip orientation and the physical surface slant:

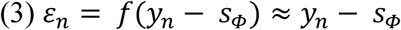

Note that in Equation 3 we have opted to approximate the actual error signal by the difference between the planned grip orientation and the physical slant. This is not meant as a mechanistic claim that these two quantities are directly compared by the nervous system; in reality, this difference is simply the source of other error signals, such as discrepancies in the expected and actual timing or magnitude of contact forces (*i.e.,* sensory-prediction errors; [18]). Having detected an error, the system updates the state of the visuomotor mapping for the next trial *x_n+1_* by extracting some of the error *ε_n_* according to the learning rate *b*:

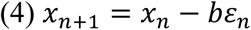

So, with repeated reaches, error signals trigger cumulative adjustments to the visuomotor mapping, without necessarily modifying perception: planned grip orientations approach the physical slant while the slant percept remains stable. If the conflict between stereo and texture is removed, then reaches should still be biased by the now-adapted internal state and give rise to an aftereffect, often considered the hallmark of adaptation.

This model of sensorimotor adaptation to a biased depth cue can account for some dissociations between perceptual judgments and visually guided actions [5] without supposing a fixed preference for stereo information in motor planning. Rather, the model maintains that motor planning relies on the same 3D shape estimates as perception, but is additionally shaped by the contributions of an adjustable internal state. This idea was the focus of Experiment 1, where we examined adaptation of the grip orientation during repeated Grip Placement (Fig 1c) on three surface slants defined by texture and stereo cues, with haptic feedback that matched the texture cue but was consistently 30° deeper than the stereo cue (*i.e.*, the stereo cue was biased).

We tested two specific hypotheses in Experiment 1: (1) motor planning relies on perceived slant, without regard for the specific mixture of slant cues, and (2) when perceived slant is biased by a faulty cue, the resulting movement errors cause proportional adjustments of the planned motor output on subsequent trials. To set up a straightforward test of the first hypothesis, we began the experiment with a perceptual Matching task, asking participants (N = 15) to produce two sets of stimuli that were perceived to have the same slants, but were composed of different combinations of stereo and texture information. This was done by adjusting the slant of a cue-consistent stimulus (*s_T_* = *s_S_*; yellow line in Figs 2a and 2b) in order to match each of three fixed cue-conflicts (*s_T_ = s_S_ + 30°*; blue and red lines indicating texture and stereo slants in Figs 2a and 2b). These 3D perceptual matches are sometimes called *depth metamers*: objects perceived to have the same depth, but with different combinations of the available depth cues (the name borrows from the psychophysical phenomenon of color metamerism, where different spectral distributions can elicit the same perceived color). Having obtained these perceptual matches, we predicted that equivalent motor responses would be produced when we suddenly switched from a cue-consistent stimulus to its matched cue-conflict, or vice versa, in the Grip Placement task.

**Fig 2.**
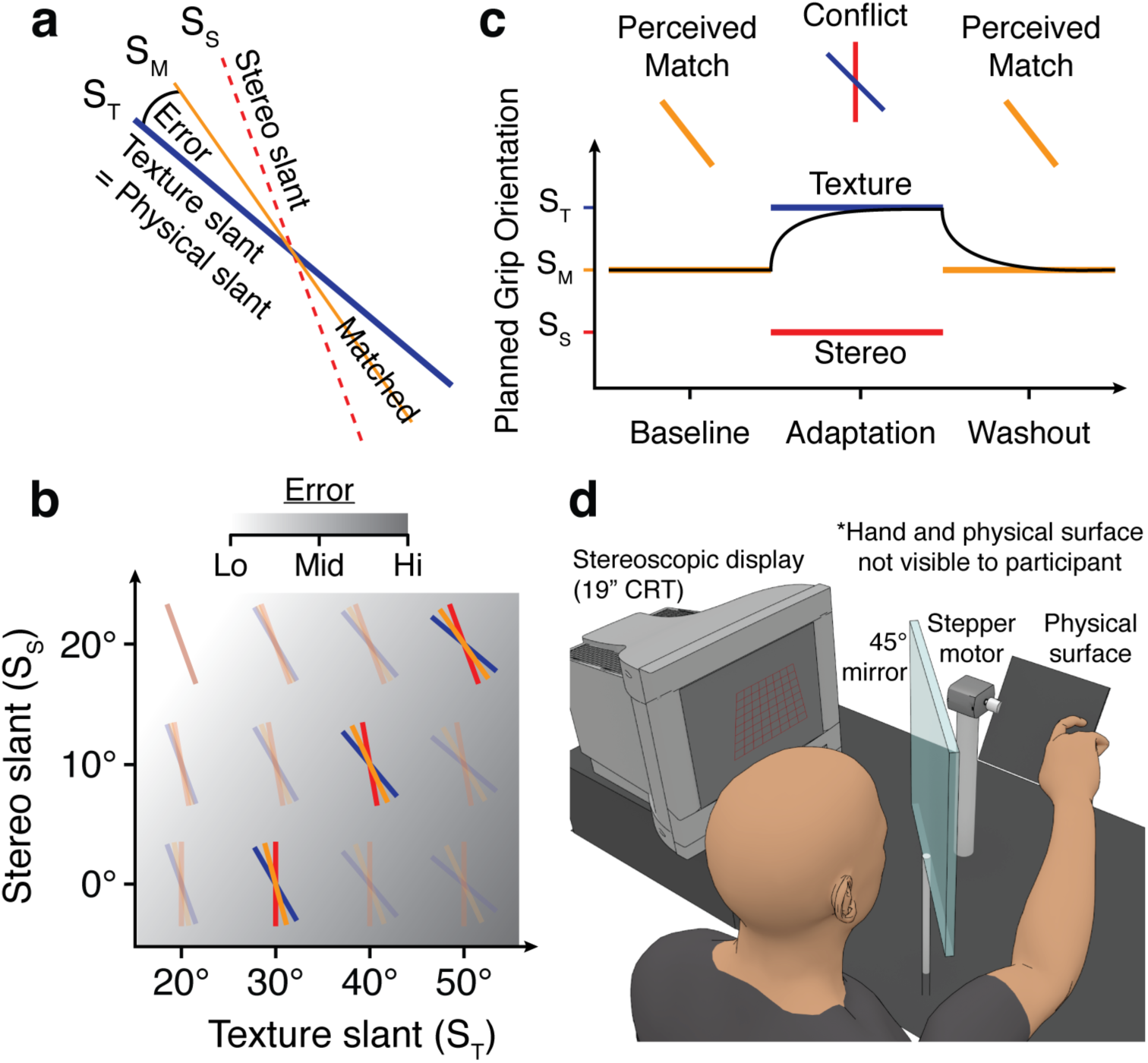
Experiment 1: Adaptation to a constant bias in stereo. (a) A cue-conflict surface. Texture (blue) specifies the physical slant, while stereo (red) shows an underestimation bias; the perceived slant (yellow) is in between. Aiming at the perceived surface will produce a movement error. (b) The three highlighted stimuli were used in Experiment 1. Stereo was consistently 30° shallower than texture slant, so errors were relatively constant (diagonal orientation of background gradient). (c) Timeline of Grip Placement task. In Baseline, cue-consistent surfaces (yellow) were presented to establish normal movement coordination (black trace). In Adaptation, cue-consistent surfaces were replaced by perceptually matched cue-conflicts (blue/red). On the first trial of Adaptation, we predict the planned grip orientation (black trace) will be identical to Baseline, mirroring the perceptual equivalence of the stimuli. Planned grip orientations should then adapt toward the physical slants, which match the texture slants. At the transition to Washout, we switch back to cue-consistent surfaces, again predicting no sudden change in the motor response due to perceptual equivalence. Thereafter, we predict rapid convergence on Baseline performance. (d) The multisensory virtual reality rig. The participant reaches with the right hand in a precision grip, orienting the hand in order to place index finger and thumb simultaneously on the observed surface, with haptic feedback from a physical surface aligned with the visual stimulus.

After the Matching task, participants completed the Grip Placement task, which followed a standard, three-phase adaptation design (Fig 2c). Participants reached forward with a precision grip, controlling the grip orientation so that index finger and thumb would contact the surface simultaneously (Fig 2d). In the Baseline phase (Fig 2c, left), each participant interacted with the personalized set of cue-consistent stimuli they indicated during the Matching task. This phase emulated well-calibrated visuomotor coordination—the physical slant encountered at the end of each movement matched both cues. At the transition to the Adaptation phase (Fig 2c, middle), the three cue-consistent stimuli were suddenly replaced by the three fixed cue-conflicts (Fig 2b). Here, we predicted that the grip orientation would not suddenly change because the stimuli were perceptually matched—notice how the black curve in Fig 2c is at the same level as Baseline on the first Adaptation trial. In contrast, if the relative weight of stereo were greater in visuomotor than perceptual tasks, we would expect the motor response to shift downward on this first trial, following the change in stereo slant. Thereafter, the physical slants reinforced texture, so we predicted that the grip orientation would rapidly shift upward toward the texture slants. Finally, we switched back to the cue-consistent surfaces in a Washout phase (Fig 2c, right); once again, we predicted no sudden change across the transition due to the perceptual matching, followed by rapid convergence on Baseline performance.

#### Perceptual Matching task

The results of the Matching task are presented in Fig 3a, with mean cue-consistent slants of 15.6°, 28.8°, and 39.3° (yellow) for the three fixed cue-conflicts (*s_S_*/*s_T_ =* 0°/30°, 10°/40°, 20°/50°; red/blue). These data correspond to a relative weight on texture information *w_T_* of 0.60 (SEM = 0.08), according to *smatch = w_T_s_T_ + (1 − w_T_)s_S_*. We computed a relative weighting of texture and stereo information here, rather than a freely varying slope parameter, because the Matching task does not provide an estimate of absolute perceived slant; it only indicates the relative influences of the two conflicting cues (as discussed by [19]). The relative weight of texture was relatively constant in the tested range, though there was a slight trend toward increased influence at greater slants, consistent with previous work [20, 21].

**Fig 3.**
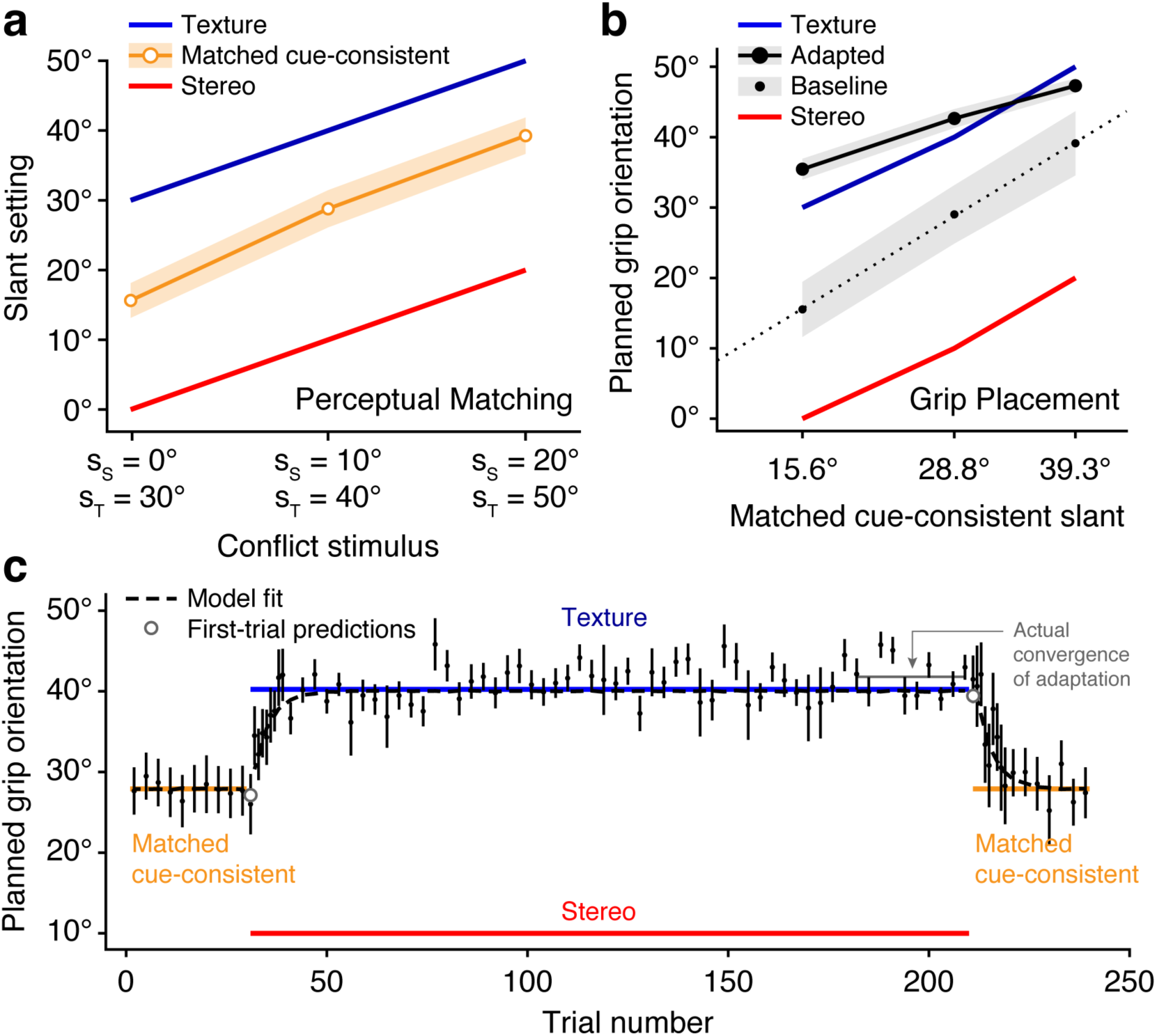
Experiment 1 results. (a) Matching: Cue-consistent slant settings (yellow) were between the component stereo (red) and texture (blue) slants of the cue-conflict stimuli. (b) Grip Placement: Average grip orientations as a function of perceived slant, in Baseline and during the final 30 trials of Adaptation (“Adapted”). The dependent variable from panel a is plotted on the x-axis; this is why the solid red and blue lines (stereo and texture slants) are not straight. In Baseline, we assume the planned grip orientations were the physical slant values (see Methods for details); this is why these data points (small points) fall on the (dotted) unity line. By the end of Adaptation (large points), planned grip orientations shifted toward the texture slants, which were reinforced by haptic feedback. (c) Grip Placement: Timeline of planned grip orientations, averaged over the three target surfaces. Our model accurately predicted the absence of any sudden change in grip orientation in the first trials of Adaptation and Washout (gray dots), and the time course of adaptation (dashed black line) was closely fit with an error-correction rate of 0.21. There was a slight underestimation of the actual convergence point of adaptation (compare model fit with gray line in trials 181 - 210; see also panel b, Texture versus Adapted)—we manually adjusted the internal state of the model on the first trial of Washout to account for this discrepancy.

#### Grip Placement task

The results of the Grip Placement task are presented in Figs 3b and 3c. Fig 3b depicts the average planned grip orientations for each of the three cue-consistent match slants during Baseline (small circles), where the physical slants matched the perceived slants, as well as during the final 30 trials of Adaptation (large circles), after exposure to the perceptually matched cue-conflicts where haptic feedback was deeper than the stereo slant, but consistent with texture. We found that the planned grip orientations from the final 30 trials of Adaptation closely matched the physical slants, though on average slightly exceeding them.

Most importantly, the timeline of planned grip orientations depicted in Fig 3c (averaged over the three targets) is highly consistent with our model of adaptation to a constant bias. At the transition from Baseline to Adaptation, where the stereo slant decreased considerably, the average grip orientation did not change (p = 0.55). If visuomotor behaviors were more sensitive to stereo information than perception, as posited the two-streams hypothesis, the grip orientation would have shifted to follow the change in stereo slant. To illustrate, suppose the perceptual weights were equal, *w_T_* = *w_S_* = 0.5. In this case, the switch from a cue-consistent slant of 25° to its perceptually matched cue-conflict would involve increasing the texture slant and decreasing the stereo slant by the same amount, say ±15°, yielding a texture slant of 40° and a stereo slant of 10°. Now, if we assume visuomotor weights that favor stereo, say *w_T_* = 0.25 and *w_S_* = 0.75, then this cue-conflict stimulus would produce a planned grip orientation of 40° × 0.25 + 10° × 0.75 = 17.5°, closer to the stereo slant than to the perceived slant. Therefore, the lack of any change on the first trial of Adaptation is evidence of a common cue-combination function in perception and action. Following this initial trial, grip orientations rapidly shifted toward the reinforced texture slants. On the first trial of Washout, the planned grip orientation again matched the model prediction almost perfectly, with no sudden shift following the change in stereo slant. Thereafter, planned grip orientations converged back to their Baseline values in response to haptic feedback from the now-shallower physical slants. Model fitting to the complete time series estimated an error-correction parameter *b* of 0.21, indicating the rapid rate of learning achieved by sensorimotor adaptation.

### Experiment 2: Cue reweighting reduces the influence of an unreliable depth cue

Experiment 1 shows that the visuomotor preference for stereo information reported in previous studies with a biased texture cue could be the result of averaging over a rapid sensorimotor adaptation process. However, this argument can only be applied to situations involving an approximately constant bias in the faulty cue, as sensorimotor adaptation is limited to uniform shifts of the motor output (assuming no anatomical [22, 23], spatial [24], kinematic [25, 26], or other features are available to consolidate adaptation within specific error contexts). Thus, for our account to be comprehensive, we must also explain how a preference for stereo information arises when other cues are unreliable, which leads to variable errors across a set of objects, as in [6].

The adaptation process modeled in Experiment 1 assumes that adjustments of the motor output can only be applied on top of the combined 3D shape estimate *ŝ = Ψ*(*s_T_, s_S_*). Yet previous work suggests it is possible to separately modify the influence of each cue prior to their perceptual combination, a process termed cue reweighting [12–15]. In the absence of sensory feedback, the relative weights of depth cues depend on a variety of factors, generally trading off in a way that favors cues with the greatest sensitivity to physical depth in the current viewing context [19–21, 27]. Here, however, we are concerned with gradual changes in cue weights that are driven by sensory feedback obtained during visuomotor interactions. To illustrate, consider a set of physical surfaces with varying slants. Both texture and stereo cues are available, but the texture cue is unexpectedly noisy, perhaps due to an irregular pattern of surface markings. Texture slant signals will therefore show a poor correlation with physical slant. As a result, visuomotor interactions with these surfaces will generate both positive and negative errors in haptic feedback, depending on whether this unreliable cue has indicated a spuriously large or spuriously small value. Since there is not a constant bias, sensorimotor adaptation (as modeled above) will fail due to interference between opposite error corrections. In contrast, cue reweighting changes the influence of each cue by adjusting cue-specific gains, making it well-suited to reduce the variable errors arising from an unreliable cue. To capture this, we can rewrite the visuomotor mapping in Equation 2 as a linear function of the individual cues *s_T_* and *s_S_*:

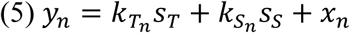

In this model, cue reweighting involves tuning the slope coefficients *k_T_n__* and *k_S_n__* in order to reduce the influence of unreliable cues and increase the influence of reliable ones. Note that when *y_n_* is a kinematic measure of the visuomotor response, as in our analysis, estimates of the slopes obtained by linear regression capture the combined effects of multiple processes: (1) the slope that relates the simulated slant values in the stimulus to single-cue slant estimates (the “single-cue mapping”), (2) the slope that captures the relative weight of each cue in the combination process (the “cue weight”), and (3) the slope that relates the cue-combined slant estimate to our measurement of the motor response (the sensitivity of the kinematic landmark). However, in an additive linear model, these various slopes simply multiply together, so we have opted to estimate and present the slope coefficients in the compact form of Equation 5.

To see how slope changes could arise through an error-based learning mechanism, consider the pattern of error signals that occurs across different values of an unreliable cue: spurious high values misleadingly increase the motor output, causing larger error signals, whereas spurious low values will decrease the motor output, causing smaller error signals. In other words, error signals will be positively correlated with the values of an unreliable cue. This fact can be exploited to perform slope adjustments with a simple rule for online supervised learning:

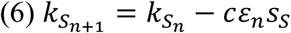

where *c* is a small, positive learning rate, *s_S_* is the input from a given depth cue (in this case, subscript *S* denotes stereo), and *k_S_n__* is the associated slope parameter on trial *n*. Through simulation, one finds that this learning rule is most robust when paired with rapid adjustments that compensate for constant errors, as modeled in the previous section. When constant error is removed, variable errors due to the unreliable cue will be centered on zero, such that spurious small values of this cue cause negative errors, yielding a small negative product in the second term (prior to the subtraction), and spurious large values of this cue cause positive errors, yielding a large positive product in the second term. Therefore, if errors are centered, on average the second term will be positive for an unreliable cue and the associated slope will be gradually reduced. Notice that under a positive constant error, the product in the second term yields even larger positive values for an unreliable cue. Although this might seem desirable because it would more rapidly reduce the influence of this cue, positive constant error actually produces inappropriate reductions in the slopes associated with *all* available cues, including those that are most reliable. Negative constant error, on the other hand, initially causes the influence of the unreliable cue to *increase*, until even more dramatic increases in the slopes associated with reliable cues cause the unreliable cue’s slope to be driven back down. In sum, the unstable behavior of this learning rule under constant error suggests a sensible complementarity with the simultaneous process of sensorimotor adaptation. Intriguingly, these observations also show why this learning rule predicts that interfering error signals are necessary to elicit cue reweighting, consistent with our previous findings on this topic [15]; we will revisit this prediction in Experiment 3B.

To examine cue reweighting in a natural visuomotor task, in Experiment 2 we asked participants to interact with a set of stimuli where one cue is uncorrelated with haptic feedback. The target stimuli were nine different surfaces rendered with independently varying stereo and texture slants (*s_T_* ∈ {15°, 30°, 45°} × *s_S_* ∈ {15°, 30°, 45°}), in either a haptic-for-texture condition (N = 28) or a haptic-for-stereo condition (N = 20). Fig 4a illustrates the stimulus set: the three stimuli located along the identity line are cue-consistent slants, whereas the other six stimuli (off-diagonal in Fig 4a) are rendered with different degrees of cue conflict. Critically, stimuli on the opposite sides of the identity line bias perception in opposite directions with respect to haptic feedback, leading to conflicting error signals from one trial to the next. In this stimulus set, since the faulty cue is completely uncorrelated with haptic feedback, the optimal solution is to eliminate that cue’s influence and to increase the influence of the reinforced cue to match Baseline performance.

**Fig 4.**
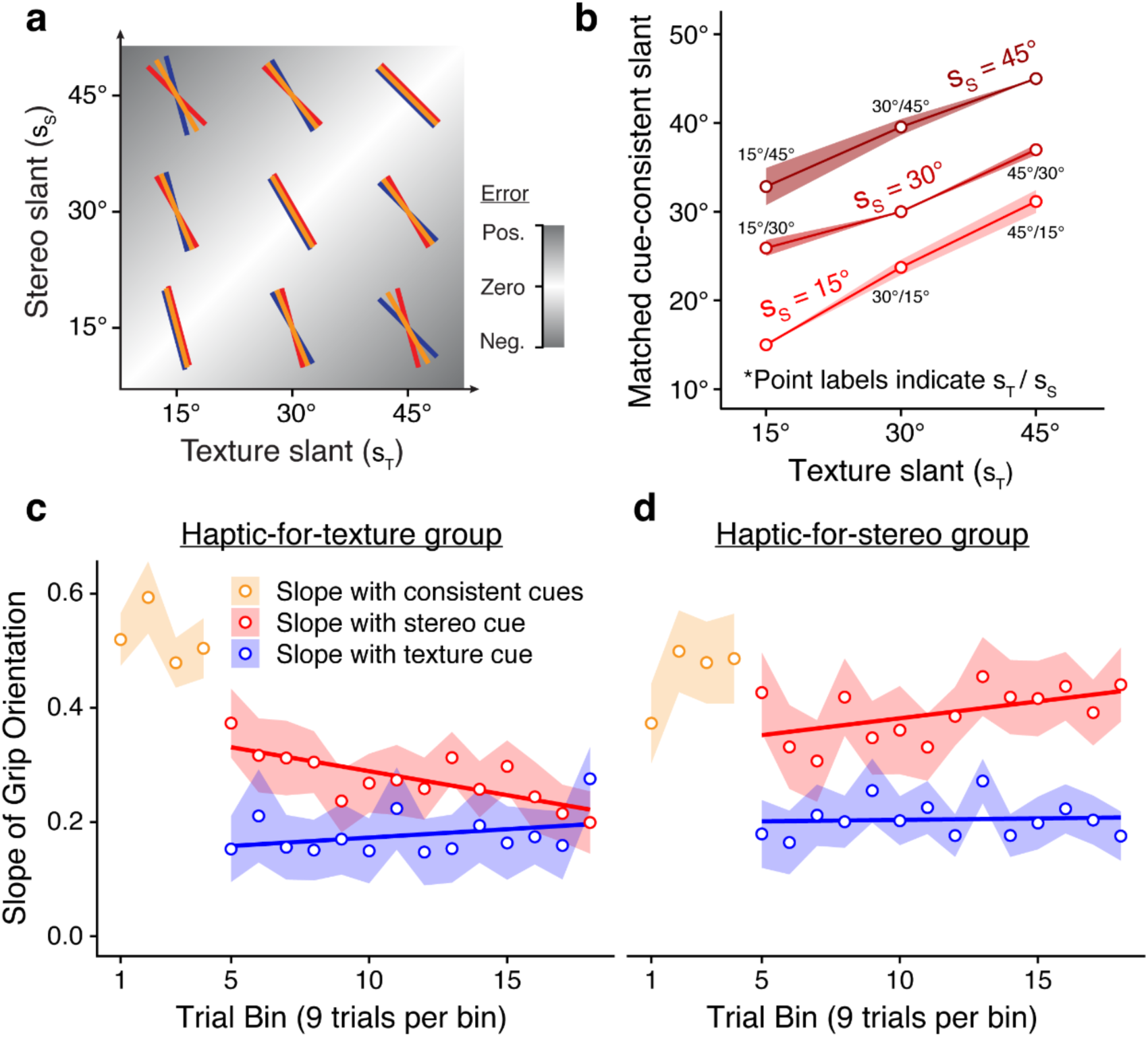
Experiment 2: Uncorrelated stimulus set and results. (a) Nine surfaces obtained by combining three texture slants (blue) and three stereo slants (red). Main diagonal contains three cue-consistent slants; the six off-diagonals are cue-conflicts. For half of the cue-conflicts (bottom-right), the expected cue-consistent match (yellow) is shallower than texture and deeper than stereo, and vice versa for the other half (top-left). As a result, when haptic feedback reinforces one cue, conflicting positive and negative movement errors should be experienced. To reduce these errors, cue reweighting is required. (b) Matching results for the nine target stimuli. Texture slant indicated on the x-axis and stereo slant indicated by the groupings (different shades of red). Perceptually matched cue-consistent slants (y-axis) were midway between the two component values of the six cue-conflicts. (c, d) Grip Placement results: In each bin, participants aimed at each stimulus once. In Baseline, we regressed grip orientations against the cue-consistent match slants (yellow). In Adaptation, we presented the nine target stimuli with physical slant reinforcing texture (panel c) or stereo (panel d). Stereo (red) and texture (blue) slopes estimated via multiple linear regression. Solid lines depict linear regression on these slopes as a function of bin number.

#### Perceptual Matching task

As in Experiment 1, participants first performed a perceptual Matching task, indicating the cue-consistent slant that appeared to match each cue-conflict in the uncorrelated stimulus set. The perceptually matched cue-consistent slants were set, on average, about halfway between the component stereo and texture slants of the six cue-conflict stimuli (Fig 4b). These data correspond to a texture weight of 0.56 (SEM = 0.04), similar to the relative weight on texture information of 0.60 found in Experiment 1.

#### Grip Placement task

Following the Matching task, participants performed the Grip Placement task. The slope coefficients estimated in each bin of this task are depicted in Figs 4c (haptic-for-texture group) and 4d (haptic-for-stereo group). During the Baseline phase, terminal grip orientations were related to cue-consistent slants by a slope of 0.50 (SEM = 0.03; yellow points), comparable to the slope of 0.60 found in the Baseline phase of Experiment 1. In the Adaptation phase, we obtained independent slopes for stereo (red points) and texture (blue points) in each bin via multiple linear regression; these indicate the sensitivity of the terminal grip orientation to each cue. In the first bin of Adaption, these slopes sum to 0.56 (SEM = 0.05), not significantly different than the mean slope observed in Baseline (p = 0.17), demonstrating that the overall sensitivity of the motor response remained the same when we changed the stimuli. Neither the Baseline slope nor the summed slopes in the first bin of Adaptation differed significantly between feedback conditions. To measure changes in the cue slopes over time, we fit additional linear regressions as a function of Adaptation bin number (solid red and blue lines). The slope coefficients obtained from these regressions indicate the bin-wise rate of change in the sensitivity of the motor response to each cue. Analyzing this rate-of-change measure using a mixed-design ANOVA (Feedback Group × Depth Cue), we found a significant interaction (*F*(1, 45) = 5.13, p = 0.028), confirming that the two feedback conditions elicited opposite changes in the relative influences of stereo and texture.

Follow-up analyses revealed a significant difference between feedback conditions in the rate of change of the stereo slope (one-tailed, two-sample *t*-test; *t*(39.79) = 3.47, p < 0.001), but not in the rate of change of the texture slope (*t*(44.06) = 0.44, p = 0.33). Additional tests demonstrated that changes in the stereo slope were observed in both conditions, significantly decreasing in the haptic-for-texture condition (mean = −0.0084 per bin, *t*(27) = 3.09, p = 0.0023) and significantly increasing in the haptic-for-stereo condition (one-tailed *t*-test; mean = +0.0059 per bin, *t*(18) = 1.86, p = 0.040). However, it is clear from Figs 4c and 4d that the observed changes fell short of the optimal form of cue reweighting that might have been achieved. Ideally, sensitivity to the reliable cue should have increased to match (or even slightly exceed) Baseline sensitivity to cue-consistent stimuli, while sensitivity to the unreliable cue should have dropped to zero. Further research is needed to identify the constraints that produced this sub-optimal cue reweighting.

Additionally, similar to Experiment 1, we found evidence that on the first trial of Adaptation, in their first interaction with a cue-conflict object, participants’ terminal grip orientations were extremely well predicted by their average grip orientation during Baseline during interactions with the cue-consistent perceptually matched object (Fig 5; Pearson’s r = 0.61, *t*(45) = 5.20, p < 0.0001). These single-trial data are noisy, but critically we find that the changes in grip orientation from Baseline to first Adaptation trial (*i.e.,* the residuals from the black dashed unity line in Fig 5) are not significantly correlated with the change in stereo slant (Pearson’s r = 0.16, *t*(45) = 1.09, p = 0.28). This first-trial analysis is consistent with our demonstration in Experiment 1 that, prior to giving informative sensory feedback, perceptually matched slants with different combinations of stereo and texture information are treated as equivalent by the visuomotor system.

**Fig 5.**
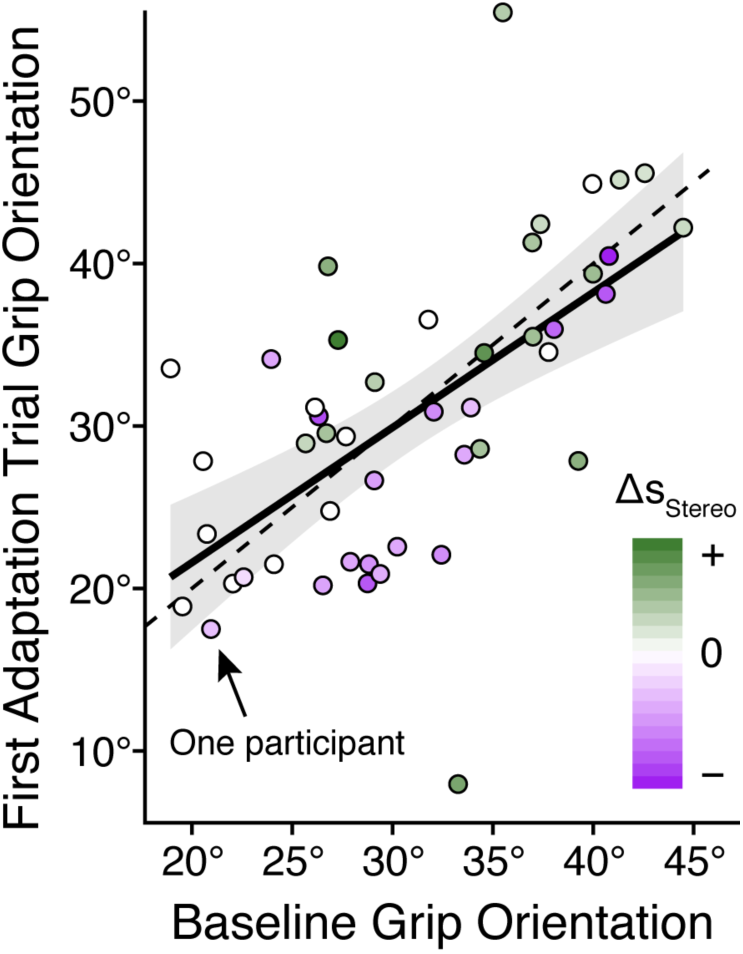
Experiment 2: Grip orientations for perceptually matched objects during Baseline and on the first trial of Adaptation. Across participants, the grip orientation on the first trial of Adaptation (the very first presentation of a cue-conflict slant) was strongly predicted by the average grip orientation of the Baseline reaches toward the corresponding matched cue-consistent slant. In contrast, the change in grip orientation at this transition (residual of each point from the dashed unity line) was not significantly correlated with the change in stereo slant from the cue-consistent slant to its perceptually matched cue-conflict slant (*Δs_S_tereo* coded by the color gradient).

To summarize, we found that the sensitivity of the terminal grip orientation to stereo information was enhanced over time when stereo was reliably correlated with physical surface slant, and reduced over time when it was uncorrelated with physical slant. Meanwhile, the sensitivity to texture information remained relatively constant throughout Adaptation, regardless of the feedback condition. However, to avoid improper interpretation of these findings, it is important to recall our discussion of Equation 5, considering that multiple processes contribute to the compact slope estimates presented here. In particular, we cannot infer from these results that the influence of stereo increased while the influence of texture stayed the same. For example, in the haptic-for-texture condition (Fig 4c), it is entirely possible that the weight of texture information in the cue-combination process increased over time, but this was masked in our data by a simultaneous reduction in the sensitivity of our kinematic measurement to changes in the cue-combined slant estimate. Such a regression toward the mean physical slant is certainly suggested by the decrease in the sum of the two slopes. Yet it is equally plausible that our observations of so-called “reweighting” are actually changes in processing of individual cues, occurring upstream of the cue-combination process (see Discussion of Experiments 2 and 3 in [28]). Both possibilities are fully consistent with existing findings on cue reweighting. For the present argument, however, the critical observation from Experiment 2 is that the relative influences of stereo and texture information shifted over time to favor the reinforced cue. The data therefore support our main claim that estimates of relative cue weights can be modified in a single session of a natural, goal-directed visuomotor task, causing them to differ from those measured in separate perceptual tasks.

Since the slope coefficients reported for Experiment 2 reflect scaling of stereo and texture signals in upstream perceptual as well as downstream motor processes, we are unable to pinpoint the locus of cue reweighting from these data. As noted in the Introduction, several previous studies suggest that cue reweighting is an inherently perceptual phenomenon that involves modification of the cue-combination function [12–14]. On this view, our hypothesis that perception and action are closely linked, rather than dissociated functions, makes the prediction that individual changes in the motor response should be correlated with concurrent perceptual changes. However, an alternative possibility, in line with the idea that perception and action rely on distinct cue-combination functions, is that cue reweighting occurs independently in visuomotor and perceptual systems, depending on the current task. With simple goal-directed actions, reweighting may be limited to visuomotor transformations, whereas tasks that involve explicit intermodal comparisons during object exploration may trigger perceptual reweighting. Most previous studies on perceptual reweighting have used tasks that fall into the latter category. To address this question, we designed a third experiment, making a few changes in our experimental design aimed at maximizing our chances of detecting potentially small changes in perception and to aid in direct comparison of perceptual and motor effects. Additionally, since we had already found cue reweighting in visuomotor interactions with slanted surfaces, and perceptual reweighting has been previously shown with nearly identical stimuli [12], we took this opportunity to investigate whether these effects would be found during grasping movements, which are based on estimates of object depth rather than surface slant. Unlike Atkins and colleagues [13], who found strong perceptual cue reweighting of scrolling-motion and texture cues when participants were asked to explicitly compare haptic and visual information while exploring an object with a precision grip, this experiment maintains our focus on natural goal-directed actions toward objects defined by stereo and texture cues. As mentioned above, showing that normal, skilled interactions are sufficient to drive cue reweighting is an important element of our argument that there is a common cue-combination function supporting both perception and action.

### Experiment 3A: Cue reweighting produces correlated perceptual and motor effects

In Experiment 3A, we asked participants (N = 36) to use a precision grip to grasp small ridge-like objects so the thumb landed on the tip and the index finger landed on a flat rear surface (Fig 6b). Thus, the relevant spatial property in Experiment 3 was object depth, as opposed to surface slant in Experiments 1 and 2, and we analyzed maximum grip apertures (MGAs, a standard kinematic measure of grasping) to quantify visuomotor cue reweighting. The switch from Grip Placement to Grasping has the added benefit of countering the argument that correlated perceptual and motor responses are found only because the motor task involves conscious, perceptually-driven planning, as opposed to the purported “vision-for-action” system recruited when performing for skilled movements like grasping [7].

**Fig 6.**
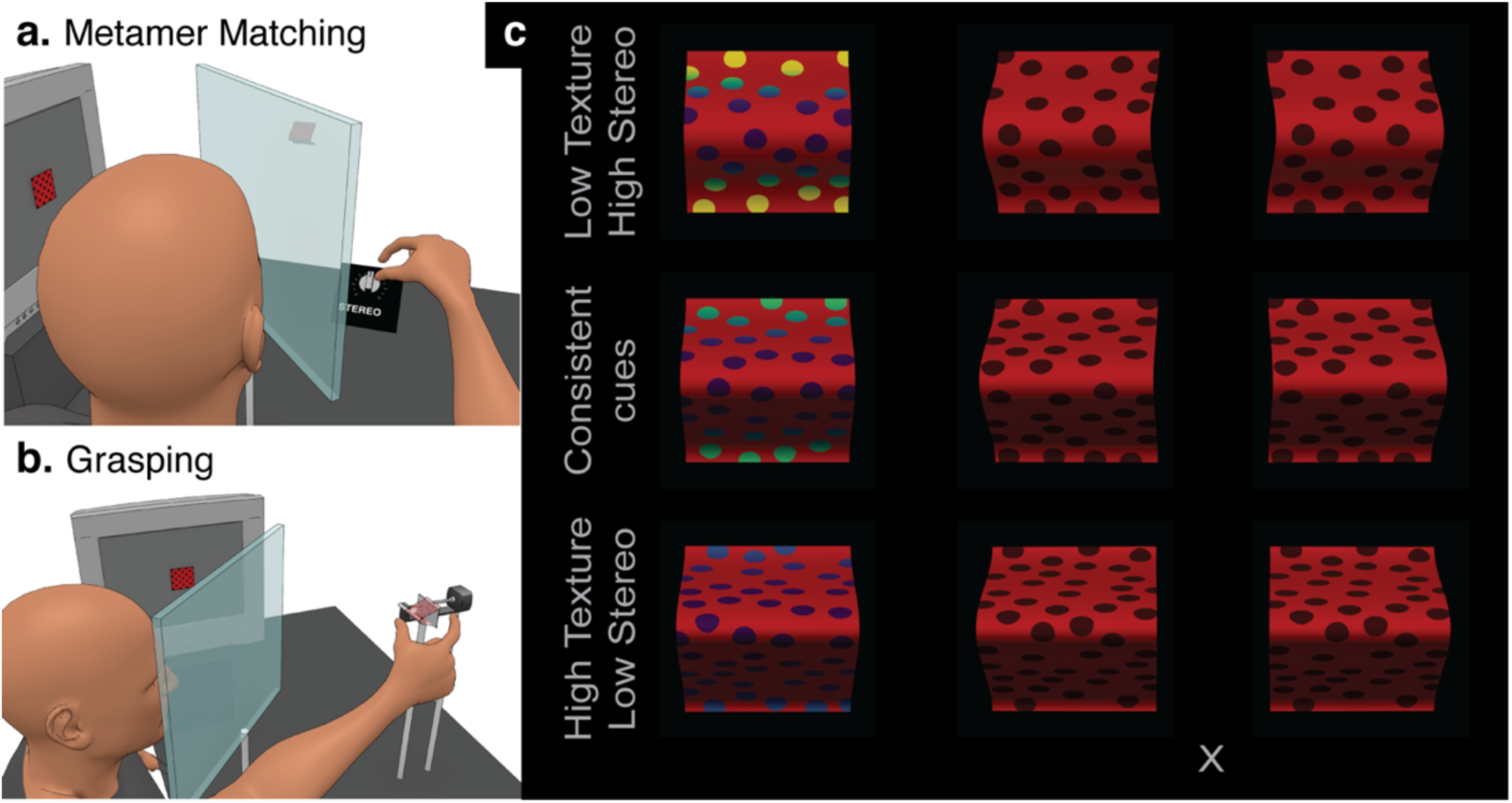
Experiment 3: tasks and depth-metamer stimulus set. (a) In the perceptual Metamer Matching task, participants created depth metamers by adjusting only the *s_T_ereo* depth (see labeled dial) of a cue-conflict stimulus to match the perceived depth of a cue-consistent standard. The adjustable cue-conflict had a fixed value of texture depth that differed from that of the standard. (b) When grasping their set of metamers, participants received haptic feedback that reinforced one of the two cues. (c) Example metamers rendered using average Pre-test results: the 30-mm cue-consistent standard (middle) was perceived to have the same depth as the combination of 18 mm of texture depth with 37 mm of stereo depth (top), as well as 42 mm of texture depth with 23 mm of stereo depth (bottom). At left, cyclopean views with stereo depth coded by a color gradient. At right, corresponding stereograms (cross-fuse).

Second, we modified the perceptual Matching task to achieve greater sensitivity to small changes in the relative influences of stereo and texture. Instead of adjusting cue-consistent stimuli to create perceptual equivalence with a fixed set of cue-conflicts, as in Experiments 1 and 2, participants in Experiment 3 began by creating a personalized set of five stimuli that were *all* perceptually equivalent to one another, based on a 30-mm cue-consistent standard. Specifically, four comparison objects were rendered with fixed texture depths (simulating 16, 24, 36, and 42 mm), and participants made incremental adjustments to the accompanying stereo depth setting (Fig 6a) until each appeared to have the same depth as the 30-mm cue-consistent standard. This personalized set of objects thus consisted of five depth metamers (*i.e.*, objects perceived to have the same depth, but with different combinations of the available depth cues).

Perceptual equivalence occurs across the set of depth metamers because the two available depth cues are inversely varied—the five increasing values of texture depth are paired with five decreasing values of stereo depth (Fig 6c shows three approximate example metamers). When the values are set according to an observer’s personal cue-combination function, they will perceive the same depth despite the differences in the component cue values. Notice that the set of depth metamers is unlike the uncorrelated set of cue-conflicts used in Experiment 2 because texture and stereo information are not manipulated orthogonally. However, it shares the key feature that when haptic feedback reinforces one of the two cues (and is inconsistent with the other), variable errors are elicited across the set. Just as in Experiment 2, these variable errors should cause interference in sensorimotor adaptation, thus requiring cue reweighting to reduce the influence of the faulty cue (which is now inversely correlated, rather than uncorrelated, with physical depth).

In this experiment, visuomotor cue reweighting can be measured as a change in the slope of the planned grip aperture with respect to the five fixed values of texture depth. According to our hypothesis, since the stimuli are initially perceptually equivalent, the planned grip apertures should also be equivalent, so the slope of the MGA across the five texture depths should be near zero at the beginning of the Grasping task. Thereafter, we predict a gradual increase in this slope measure in the haptic-for-texture condition and a gradual decrease in the haptic-for-stereo condition, as MGAs begin to follow the reinforced cue. Remember the two cues are inversely correlated, so a negative slope of the MGA with texture implies a positive slope with stereo.

Conveniently, the metamer Matching task also allowed us to compute the relative influences of the two cues in perception, so we could measure perceptual cue reweighting by having participants complete the Matching task again in a Post-test, immediately after the Grasping task. In fact, the change in perceptual weights from Pre-test to Post-test can be converted directly into predictions of the MGA slope across the metamer set (see Methods). Our main prediction for this experiment, therefore, was that the change in perceptual cue weights from Pre-test to Post-test would predict, on an individual basis, how MGAs scaled across the metamers in the Grasping task.

#### Pre-test Metamer Matching task

Pre-test Matching results are depicted in Fig 7 with black closed circles and dotted lines; colored open circles and solid lines depict the Post-test results, which will be described later. As explained above, for each of the four fixed texture depths (18, 24, 36, or 42 mm), participants adjusted stereo depth until the resulting cue-conflict stimulus perceptually matched the 30-mm cue-consistent standard. In the haptic-for-texture condition, the four fixed texture depths were paired with stereo depths of 36.9, 32.1, 26.1, and 23.1 mm, respectively (SEMs: 1.1, 0.5, 0.6, 1.0 mm), corresponding to a relative texture weight of 0.270 (SEM = 0.026). In the haptic-for-stereo condition, the adjusted stereo depths were 36.5, 31.7, 26.2, and 22.6 mm (SEMs: 1.1, 0.5, 0.5, 0.9 mm), corresponding to a texture weight of 0.263 (SEM = 0.029). The nearly equivalent Pre-test settings across conditions make sense because no visuomotor training had yet been provided.

**Fig 7.**
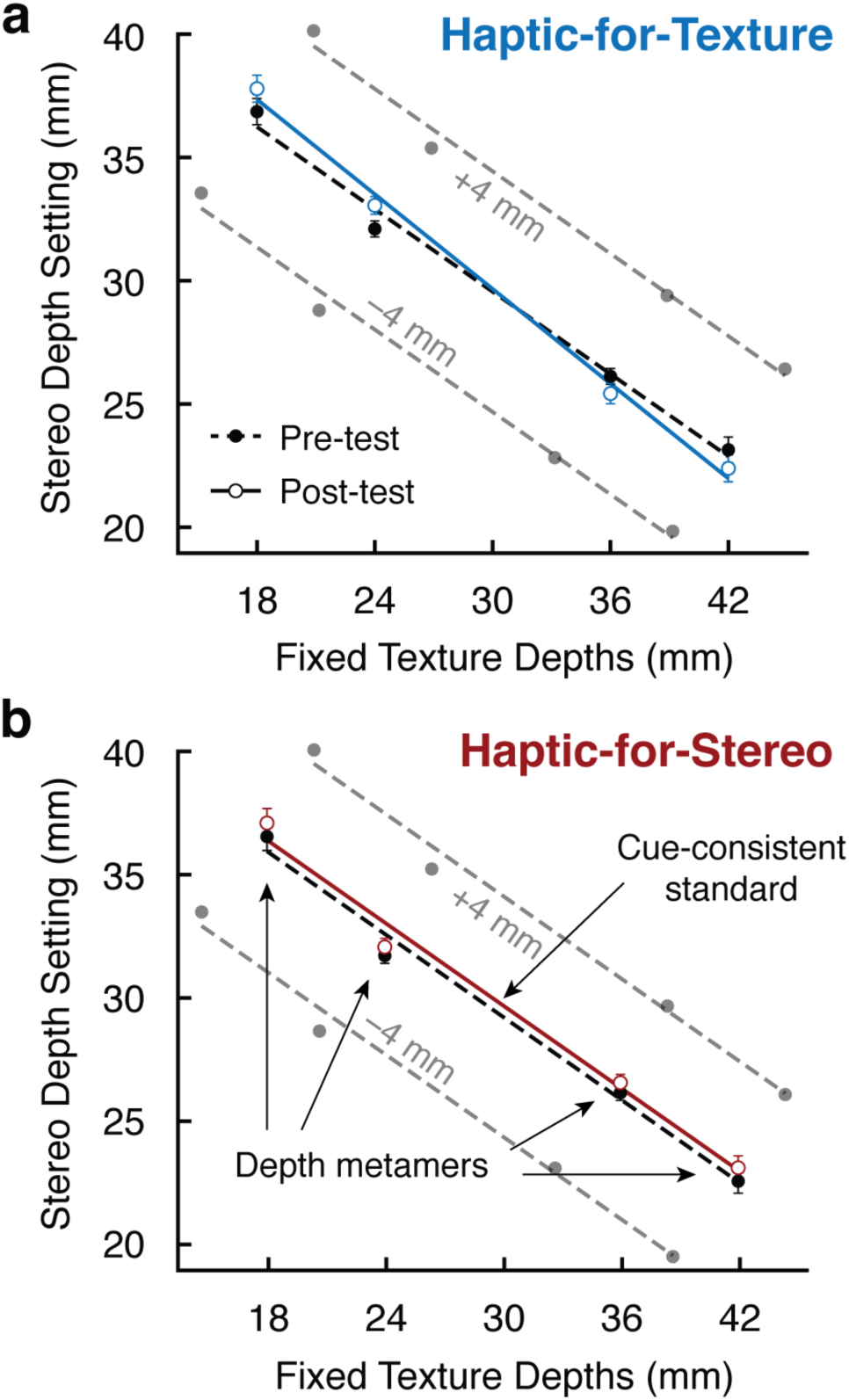
Experiment 3A perceptual results. In the Pre-test and Post-test phases (see legend), participants created a personalized set of five depth metamers. The standard was a 30-mm cue-consistent stimulus (not shown). For the other four stimuli, texture depths were fixed at 18, 24, 36, or 42 mm (x-axis) and the participant adjusted stereo depth (y-axis) until the perceived depth matched the standard. Gray points and dotted lines depict the additional stimuli presented in the Grasping task in order to provide some perceptible variability in depth. (a) In the haptic-for-texture condition, the relative influence of texture information increased from Pre-test to Post-test; stereo settings are higher at left and lower at right. These data show that after Grasp training, stronger stereo signals were required to balance out the same weak texture signals (18 and 24 mm) and make them equivalent to the 30-mm cue-consistent standard, while weaker stereo signals were required to balance out the same strong texture signals (36 and 42 mm). (b) No perceptual change was observed in the haptic-for-stereo condition; stereo settings were the same in Pre-test and Post-test.

As expected, stereo depth settings were inversely related to the fixed texture depths: the largest stereo depths were paired with the smallest texture depths, and the smallest stereo depths were paired with the largest texture depths. Thus, when presenting these stimuli in the Grasping task, haptic feedback could be positively correlated with one cue (the reinforced cue) and negatively correlated with the other (the faulty cue) in order to elicit cue reweighting. Note, however, that we opted to include additional stimuli in the Grasping task in order to provide some variation in perceived depth, aiming to maintain participants’ interest in the task. Our reasoning was that if all the stimuli were perceived to be equal in depth, participants may simply give up trying to scale their grip apertures with the objects. To provide some perceptible variability in object depth that correlated with haptic feedback, we replicated the set of five metamers specified by each participant in the Pre-test to create three distinct sets (gray circles and dotted lines in Fig 7; see Methods for details on how these were generated).

#### Grasping task

The Grasping task began with a 26-trial Baseline phase. In the first half of this phase, participants grasped a range of cue-consistent objects spanning 18–42 mm in depth; the metamers were not yet introduced. In these trials, MGAs reliably scaled with variations in cue-consistent depth (slope of 0.64, SEM = 0.05; not depicted in Fig 8).

**Fig 8.**
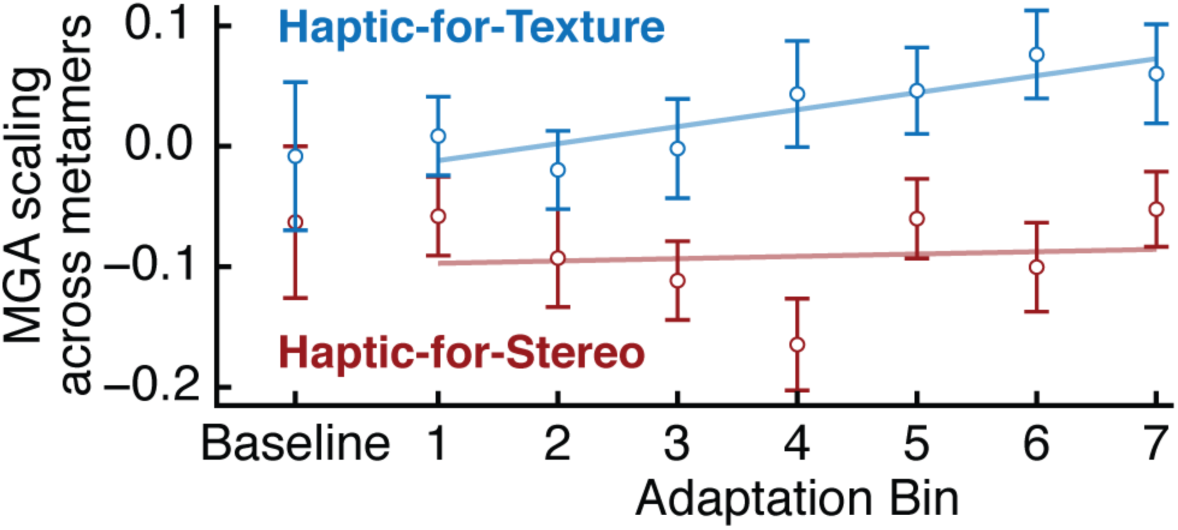
Experiment 3A Grasping results. Timeline of MGA scaling across the metamer set (slope with respect to the fixed texture depths spanning 18-42 mm). When metamers were first introduced (Baseline), MGA slopes were near zero, mirroring the perceptual equivalence. Across the Adaptation phase, positive-trending slopes were observed in the haptic-for-texture condition, indicating increased reliance on texture. However, negative-trending slopes, which would indicate increased reliance on stereo, were not observed in the haptic-for-stereo condition.

In the second half of Baseline, we maintained five cue-consistent stimuli but also introduced the five depth metamers created in the Pre-test. At this point, the haptic feedback associated with each metamer was kept at 30 mm, allowing us to measure grasp performance prior to receiving any feedback indicating that the haptic depths differed. As predicted, participants used roughly the same MGA across the metamers, mirroring the perceptual equivalence of these stimuli (Fig 8a, Baseline bin; MGA slope with respect to the fixed texture depths was −0.01 in haptic-for-texture, −0.06 in haptic-for-stereo, both SEM = 0.06). In contrast, if grasp planning depended on separate 3D shape processing with a hardwired preference for stereo information, we should have observed significantly negative slopes, since stereo depth is inversely related to texture depth, and texture depth is the predictor in our regressions. This result demonstrates that depth metamers elicit indistinguishable visuomotor responses despite their differing combinations of stereo and texture information, just as we found when switching between cue-consistent and cue-conflict slant metamers in Experiments 1 and 2.

Next, in the Adaptation phase, participants were exposed to the full set of metamers, with haptic feedback reinforcing one of the two cues. For example, in the haptic-for-texture condition, metamers with larger texture depths (and smaller stereo depths) were paired with deeper physical objects, and those with smaller texture depths (and larger stereo depths) were paired with shallower physical objects. In the haptic-for-stereo condition, the physical object sizes remained the same, but they were positively correlated with stereo and therefore negatively correlated with texture. As explained in the introduction to Experiment 3A, cue reweighting in the haptic-for-texture condition would be shown by the MGA slope with texture becoming gradually more positive, while in the haptic-for-stereo condition it would be shown by the slope becoming gradually more negative. The time course of the MGA slope with texture in each condition is shown in Fig 8a. Linear regression on the slopes as a function of bin number revealed an overall positive rate of change in the haptic-for-texture condition (*t*(35) = 2.76, p = 0.0046), indicating that participants learned to differentiate the metamers according to the haptic feedback. However, despite a noticeable trend in early Adaptation bins, the haptic-for-stereo condition failed to produce a stable decrease in MGA scaling across the metamer set (p = 0.63).

The asymmetry in cue reweighting between our two feedback conditions continues an unexpected trend that has become evident in our recent work: haptic-for-stereo conditions consistently elicit milder cue reweighting than corresponding haptic-for-texture conditions (Experiment 2, compare Figs 4c and 4d; [12, 15]). One explanation is that this reflects a ceiling effect due to the already-dominant influence of stereo information in many observers. We may have unintentionally made this worse by using metamer set, rather than fixed values of stereo and texture depth as in Experiment 2, as stereo-dominant observers now had only minimal variation in the stereo cue, thus limiting their ability to learn to use this variability as a predictor of physical depth. Indeed, the Pre-test results show that, on average, the range of stereo depths across the metamers was 14 mm, whereas the range of texture depths was 24 mm. Another possible explanation of the asymmetry arises if one considers that the error signals that drive cue reweighting could be sensory-prediction errors, *i.e.*, differences between actual sensory feedback and the expected sensory feedback, predicted during movement planning on the basis of the cue-combined perceived shape and the outgoing motor command. Sensory-prediction errors would necessarily be smaller in conditions where feedback is consistent with the dominant depth cue. In any case, additional research is warranted in determining the cause of this rather consistent asymmetry across experiments. Lastly, to avoid confusion, we should emphasize that this asymmetry in the degree of reweighting between haptic-for-texture and haptic-for-stereo conditions is not the same as the asymmetry found *within* each these conditions, where the sensitivity of the motor response to stereo changes, while sensitivity to texture remains quite stable (see Figs 4c, 4d and [15]).

#### Post-test Metamer Matching task

Returning to Fig 7, we can evaluate changes in the Matching task stereo settings from Pre-test (solid circles, dotted lines) to Post-test (open circles, solid lines) as evidence of perceptual cue reweighting. In the haptic-for-stereo condition (Fig 7b), the relative weight of texture information was 0.259 (SEM = 0.031), a small decrease from the value of 0.263 measured in the Pre-test. In the haptic-for-texture condition (Fig 7a), the Post-test texture weight was 0.308 (SEM = 0.032), up about 4% from the Pre-test value of 0.270. Most importantly, these changes in relative cue weights were found to be significantly modulated by feedback condition (*t*(35) = 1.77, p = 0.043). Follow-up *t*-tests revealed significant perceptual reweighting in the haptic-for-texture condition (*Δw_stereo_* = −0.038, *t*(35) = −2.43, p = 0.010), but not in the haptic-for-stereo condition (*Δw_stereo_* = 0.004, *t*(35) = 0.20, p = 0.42).

The main goal of this experiment was to determine whether perceptual and visuomotor cue reweighting were caused by a modification of the same cue-combination function, as opposed to being to independent types of learning. Since our haptic-for-stereo condition was ineffective in eliciting either type of cue reweighting, it is not useful for this analysis. However, since we observed both forms of cue reweighting in the haptic-for-texture condition, we can analyze this condition to determine whether these changes were correlated on an individual basis, which would suggest a common source. This analysis produced the key result of the Experiment 3: individual perceptual changes, converted into predictions of MGA scaling across the metamers during Adaptation, were significantly correlated with the measured MGA slopes (Pearson’s r = 0.49, *t*(34) = 3.30, p = 0.0023; Fig 9).

**Fig 9.**
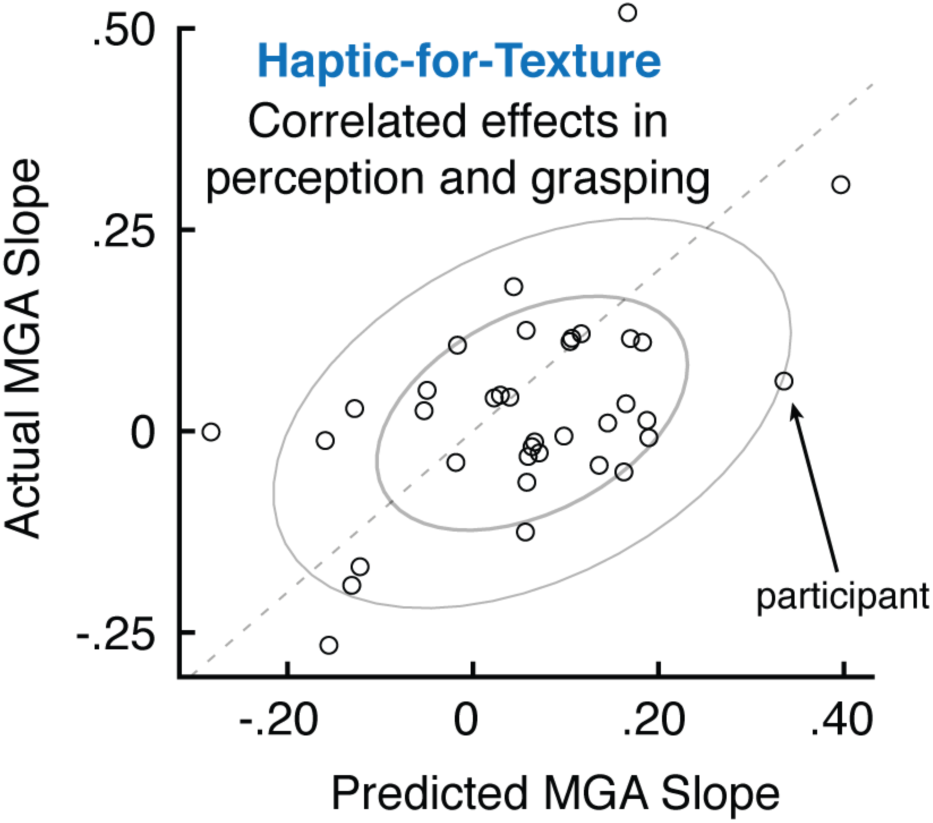
Individual perceptual changes predict grasp performance during Adaptation. In the haptic-for-texture condition of Experiment 3A, where we found motoric and perceptual changes, the predicted MGA slope across the five metamers for each participant (based on their measured perceptual change) was correlated with their actual slope.

### Experiment 3B: A biased cue induces sensorimotor adaptation without cue reweighting

If the perceptual cue reweighting observed in Experiment 3A was indeed the result of variable movement errors, as we have suggested, then perceptual changes should not occur during exposure to a biased cue, since constant movement errors can be resolved by sensorimotor adaptation (see Experiment 1). To test this prediction, we ran a control experiment with a new group of participants (N = 20). The design was identical to the haptic-for-texture condition of Experiment 3A, except instead of presenting metamers during the Adaptation phase, we presented objects with a fixed cue-conflict, as in Experiment 1: during Adaptation, stereo depths were 8 mm shallower (10–34 mm in 6-mm increments) than the physical depths, which remained consistent with texture depths (18–42 mm).

Fig 10a shows that the MGA time course from this control experiment is well captured by an exponential fit, consistent with the proportional error-correction of sensorimotor adaptation. When stereo depths decreased by 8 mm at the onset of Adaptation, the MGA suddenly dropped by 5.75 mm. This change in the motor response matches our prediction based on the measured weights of the two cues in Pre-test Metamer Matching (see prediction interval in Fig 10a, bin 4), once again indicating a common cue-combination function for perception and action. Subsequently, MGAs increased and leveled off near their Baseline values, fully compensating for the biased stereo cue. Unlike in Experiment 3A, here we found no increase in the relative weight of texture information from Pre-test (0.246) to Post-test (0.240); participants indicated nearly identical sets of metamers in both test phases (p = 0.63; Fig 10b). Moreover, the perceptual changes found in Experiment 3A were significantly greater than those observed in this experiment (Welch’s two-sample *t* test; t(41.855) = 1.78, p = 0.041; Fig 10c). This is consistent with our hypothesis that cue reweighting is driven by altered correlations with haptic feedback [15], and not simply by mismatches between the faulty cue and haptic feedback. Note that participants received approximately equivalent exposure to visual-haptic mismatches in both experiments, in terms of number of trials and average magnitude of the mismatches, but the constant errors in Experiment 3B were quickly resolved by sensorimotor adaptation, whereas the variable errors in Experiment 3A persisted throughout the training session.

**Fig 10.**
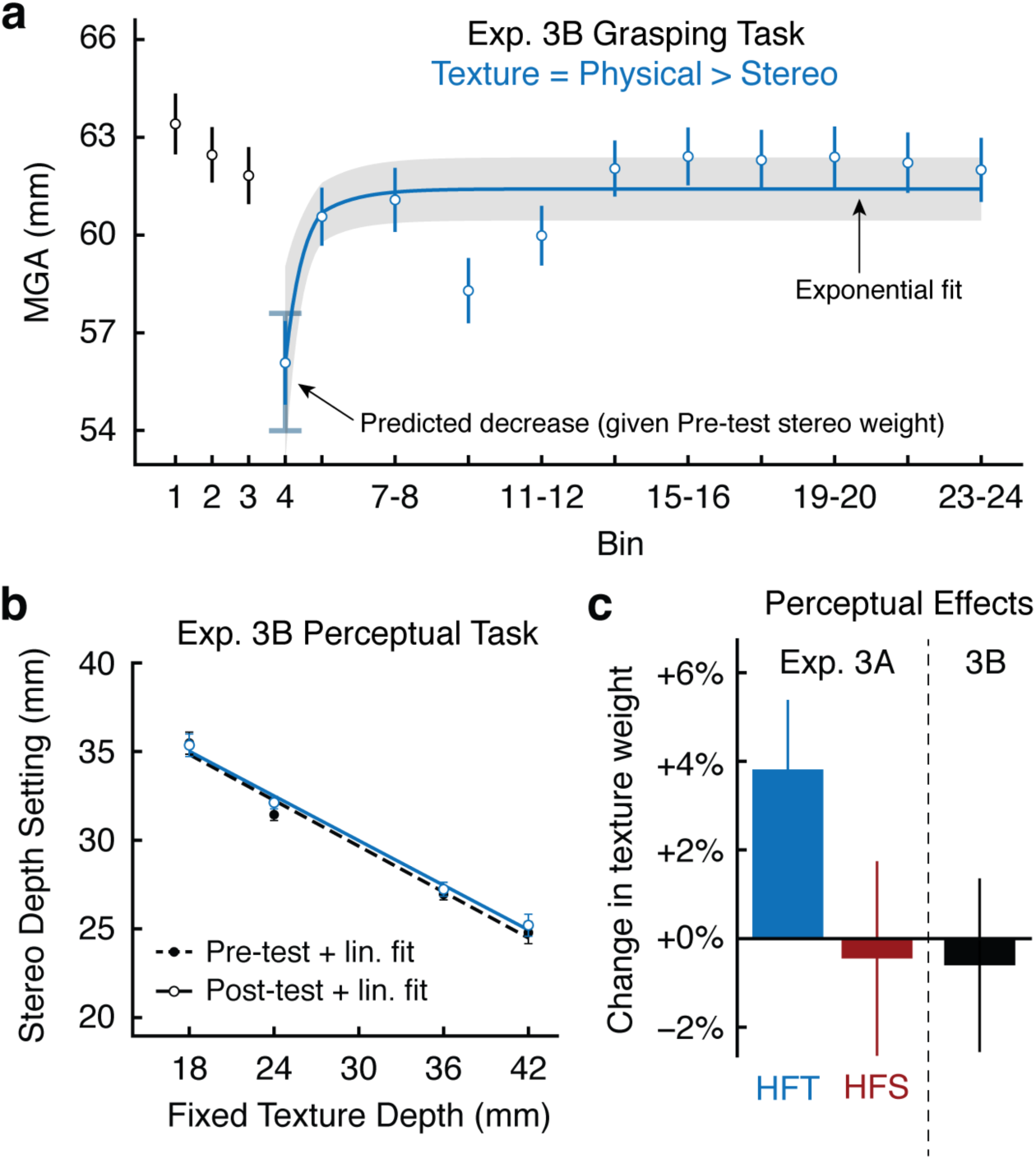
Experiment 3B results and Experiment 3 summary. (a) When stereo depth was suddenly decreased by 8 mm at the transition to from Baseline (black circles) to Adaptation (blue circles), MGAs decreased by about 6 mm, matching a prediction based on the Pre-test stereo weight of 0.75 (error bar interval in bin 4). Across the Adaptation phase, MGAs returned to Baseline levels following an exponential function (blue fit line). (b) No changes were observed in the metamer stereo settings from Pre-test to Post-test. (c) Summary of the perceptual effects from Experiments 3A and 3B.

## Discussion

In four experiments, we examined changes in the performance of visually guided reaching and grasping movements in response to haptic feedback, with the goal of understanding whether sensorimotor adaptation and cue reweighting help reduce movement errors caused by distorted 3D shape information. In particular, we focused on how these two distinct forms of supervised learning are appropriate for two different types of distortion: the presence of biased versus noisy depth cues. Notably, each of these types of distortion has been used in past experiments that aimed to test whether perceptual and visuomotor responses are mediated by separate cue-combination functions. The present series of experiments provides repeated demonstrations that these short-term learning processes are indeed active during natural goal-directed visuomotor behavior, and that when they are properly accounted for, the relative influences of stereo and texture information appear to be identical in perceptual judgments and motor responses.

### Sensorimotor adaptation to a biased depth cue

In each experiment, we found evidence that movement planning relies on the same cue-combined estimates of 3D shape as perceptual judgments, in contrast to previous claims that visuomotor tasks activate a separate cue-combination mechanism with a hardwired preference for stereo. In Experiment 1, we were able to demonstrate this only by carefully determining different combinations of stereo and texture information that were perceived to have the same depth, and suddenly switching between these depth metamers in an ongoing visuomotor task. This precise technique was necessary because of how rapidly sensorimotor adaptation occurs. Indeed, Experiments 1 and 3B verified that sensorimotor adaptation operates as expected when an available depth cue becomes biased, in line with the standard proportional error-correction model. The model-estimated error correction rate of 0.21 from Experiment 1 indicates a fast exponential time course, likely reflecting the combined contributions of implicit and explicit components of adaptation [29, 30]. Overall, these results support the argument that some perception-action dissociations arise because sensory feedback drives the motor response toward cues that are more physically accurate, even when the 3D shape estimate used for motor planning remains biased.

### Implications for cue recalibration

Beyond its implications for the debate on perception-action dissociations, our demonstration that sensorimotor adaptation compensates for movement errors caused by biased depth cues can provide novel perspectives on other forms of learning that are also geared toward resolving biases in depth cue processing. For instance, consider cue recalibration (*NB*: not cue reweighting), a gradual supervised learning process typically characterized as a uniform shift in the mapping of individual depth cues onto estimates of perceived world properties [28, 31, 32]. Since cue recalibration and sensorimotor adaptation ultimately have the same effects on motor behavior, they are in some sense redundant (maybe even conflicting) processes. From this perspective, we can ask why cue recalibration occurs at all when sensorimotor adaptation provides a more rapid solution. One possible answer is that cue recalibration serves a different computational goal than sensorimotor adaptation, perhaps responding to durable changes in the covariance structure of concurrent sensory inputs, rather than minimizing movement-related error signals. This raises a novel empirical question for future research: can cue recalibration be obtained with only passive exposure to haptic sensory signals? If not, it could mean that sensory-perceptual mappings and visual-motor mappings are *both* modified by movement-related errors, such as sensory-prediction errors [33].

Interestingly, the idea that sensory-prediction errors drive cue recalibration, as opposed to a passive sensory process that monitors the alignment of different sensory modalities, leads to a novel interpretation of findings from Adams and colleagues [34]. In that study, recalibration of shape-from-shading judgments was found to be stronger when observers received intermittent haptic or stereo feedback that conflicted with their initial percept from shading alone, compared to when conflicting stereo information was continuously available as they touched the targets. In the latter condition, there was a mismatch between the interpretation from shading and the interpretation from stereo, but the cue-combined percept would have been dominated by stereo information. As a result, the subsequent haptic feedback (which was consistent with stereo) would have matched the observer’s predictions. In the former case, sensory feedback was obtained only at specific moments of interaction, so it would have violated the observer’s expectations about what that feedback should have looked or felt like, since these predictions had to be formed on the basis of shading alone. The authors referred to this violation as an “oops factor”, where the new sensory signals forced a revision of the initial stimulus interpretation [34]. Notice, however, that it is essentially the same concept as a sensory-prediction error: a conflict between actual sensory feedback and an internal prediction of that feedback formed on the basis of earlier information. Below, in our discussion of cue reweighting, we will further unpack the idea that persistent sensory-prediction errors could be responsible for modifying 3D visual processing, in addition to their well-accepted role in sensorimotor adaptation.

### Cue reweighting requires altered correlations with haptic feedback

Consistent with previous perceptual studies [12–14], and extending our own recent findings from another visuomotor task [15], Experiments 2 and 3 showed that exposure to an unreliable depth cue produces cue reweighting. Whereas sensorimotor adaptation is well-suited for situations involving constant biases, cue reweighting is required in situations where the signals from one depth cue have become less correlated with physical reality. Consistent with this functional distinction, we have previously shown that cue reweighting of the motor response does not occur during exposure to constant biases, but only in response to reduced correlation of one available cue with haptic feedback [15]. This claim is bolstered by the findings of Experiment 3, where perceptual cue reweighting occurred when haptic feedback was positively correlated with texture and negatively correlated with stereo (Exp 3A), but not when haptic feedback remained correlated with both cues but was misaligned with stereo by a constant bias (Exp 3B). In the biased-stereo condition, sensorimotor adaptation drove the motor response toward the physical surface shape, compensating for the perceptual bias just as in Experiment 1. Thus, this study provides additional evidence against the naïve hypothesis that changes in the relative weighting of depth cues are made according to the absolute mismatch between each cue and haptic feedback. An important remaining question is why altered correlations between individual depth cues and haptic feedback are necessary to produce cue reweighting.

At present, most researchers approach the phenomenon of cue reweighting from the perspective of Bayesian cue combination (*cf.* [35, 19, 20]): statistically, the optimal way to combine multiple unbiased but noisy estimates of the same world property is to assign linear weights to the single-cue estimators based on their relative reliabilities (hence the name cue *reweighting*; [12–14]). From this perspective, it is natural to hypothesize that the mechanism supporting cue reweighting involves a direct estimate of the reliability of each cue. One way to coarsely estimate the relative reliabilities would be to monitor their correlations with haptic feedback over the course of repeated interactions. Although they will be noisy, these correlations could theoretically serve as proxies for the actual cue reliabilities, and cue weights could be set accordingly. This provides a straightforward answer to the question of why altered correlations with haptic feedback are necessary for cue reweighting. Although none of the present results conflict with this model, it is worth pointing out that there are other plausible alternatives that should be investigated. In the remainder of the Discussion, we describe one such alternative model, where altered correlations are important because they lead to persistent sensory-prediction errors. In particular, we suggest that sensory-prediction errors, widely believed to be the driving error signals causing sensorimotor adaptation, could also be responsible for the perceptual changes found in cue reweighting and recalibration.

### Persistent sensory-prediction errors and perceptual change

Above, we described previous findings [34] showing that perceptual processing of shading information is more strongly recalibrated by limited, intermittent exposure to feedback that generated sensory-predictions errors than by continuous exposure to inconsistencies between shading and combined stereo-and-haptic feedback that did not generate sensory-prediction errors. These findings suggest that sensory-prediction errors, computed with respect to the cue-combined percept, are more effective in changing perceptual processing than discrepancies between single-cue estimates. Moreover, since constant sensory-prediction errors are eliminated quickly by sensorimotor adaptation, this approach also explains why visual perceptual changes are quite rare in typical adaptation experiments. In contrast, studies that have succeeded in eliciting perceptual changes (*i.e.*, cue recalibration or cue reweighting) have consistently involved sensory-prediction errors that could not be fully resolved by adaptation. As explained earlier, all previous studies on cue reweighting [12–15] used a stimulus set where the correlation between the faulty cue and haptic feedback was reduced, thus leading to variable errors that would have caused interference in sensorimotor adaptation. Likewise, many cue-recalibration experiments have also involved variable errors. For the bump/dimple stimuli used by Adams and colleagues [31, 34], only some targets were paired with feedback that conflicted with the initial interpretation of the shading cue, and even within this small subset the direction of the errors varied: exploration of some perceived dimples ended up generating feedback consistent with a bump, and some perceived bumps generated feedback consistent with a dimple. In another experiment that investigated recalibration of depth-from-stereo [28], the training stimuli involved visual-haptic mismatches that ranged from 10-21 mm. This 11-mm range of errors would have prevented sensorimotor adaptation from fully eliminating sensory-prediction errors. In summary, the common thread across multiple studies, including the present one, is that changes in 3D shape perception occur precisely in those situations where sensory-prediction errors are highly variable, and therefore persistent.

### A network model of cue reweighting via sensory-prediction errors

Speculatively, we propose that the persistence of sensory-prediction errors allows their corrective effects to backpropagate through the visuomotor system, gradually producing changes in upstream visual areas that process and combine depth cues. In Equation 5, we posed the cue-combination process as a global function with cue-specific slope parameters that are modified during visuomotor interactions. However, sensorimotor adaptation research suggests it may be more appropriate to model the corrective effects of sensory-prediction errors using a distributed representation, where a population of locally-tuned units (“neurons”) encode relevant features of the motor response (*e.g.*, target location, movement velocity; [16, 25, 26], but see [36] for evidence against a distributed encoding of target location in the visuomotor mapping). In the case of interacting with 3D objects, we are instead concerned with *visual* features of the target, including stereo and texture information. When viewing a particular object, sensory input signals establish a pattern of activity across populations of stereo- and texture-sensitive units [37]. One possibility is that these units are tuned to different combinations of depth cue signals: some units might respond maximally when a strong stereo signal is paired with a weak texture signal, while others respond maximally to strong signals from both stereo and texture. When viewing an object with a large stereo depth and a small texture depth, nodes that are sensitive to this particular combination of visual information would respond preferentially, while nodes sensitive to other combinations are relatively silent. The present pattern of activity could then be mapped onto a particular perceived depth by (1) scaling the input activity of each node by a vector of weight parameters, which determine the influence of each node, and (2) pooling the weighted node activities. This approach to computing arbitrary functions is called local function approximation, since the output is constrained to vary smoothly with changes in the input dimensions (available depth cues), but unlike Equation 5 there are no explicit global parameters. Instead, the parameters are the node weights, and adjusting the weight of a node based on error feedback will only increase or decrease the perceived depth in a localized region of the input domain.

According to this model, cue reweighting could involve the same error-correction mechanisms that underlie sensorimotor adaptation, but at a deeper level of the visuomotor network where depth cues are encoded. Cue reweighting could thus be achieved entirely by incremental, local adjustments of the approximated mapping from depth cues to perceived depths. Since the nodes are selective for specific combinations of depth cues, sensory-prediction errors registered during goal-directed movements would most strongly affect the weights of the highly active nodes that encode the current combination of depth cues, with the corrective effect falling off in more distant regions with less active nodes. In fact, this behavior can be achieved simply by applying the update rule of Equation 6 to the node weights and activities in this distributed representation, rather than to the cue-specific slope parameters and metric shape estimates that appear in Equation 5. As local corrections accumulate, the resulting changes in perception become evident in the slope parameters estimated by multiple linear regression. In particular, the slope changes that we presented as evidence of cue reweighting would occur when node weights change in opposite directions in different regions of the input space, as when large values of an unreliable depth cue cause positive errors and small values cause negative errors. This is exactly the kind of situation that cannot be resolved by sensorimotor adaptation, thus allowing these error signals to persist and gradually modify upstream visual areas that are slower to adapt.

An interesting feature of this network model is that it offers a representational basis in which the contributions of individual depth cues can be distinguished, providing the separability needed for cue reweighting, while also accounting for mandatory fusion of depth cues in perception, a seemingly contradictory phenomenon. In their demonstration of mandatory fusion, Hillis and colleagues [38] showed that individual depth cues could not be isolated to aid perceptual discriminations between cue-consistent and cue-conflict stimuli. To explain this phenomenon with the proposed network model, notice that when a particular pattern of activity occurs across the population of units, the available depth cues have already been combined. Although the pattern of activity does indicate a specific combination of cue signals, thus providing separability, the component signals can no longer be individually accessed. Therefore, if perceived 3D shape is determined by learned weights on the presently activated units, then when two distinct combinations of depth signals activate populations with similar summed weighted activities, the resulting percepts will not be discriminable. This will hold true even if the individual cues are discriminable when presented in isolation, since isolating the cues entails a different pattern of population activity.

Of course, the network model is not without its drawbacks. One significant problem that arises is the combinatorial explosion inherent in representing all possible combinations of depth cues. However, this may be less problematic than it seems. First, consider that many depth cues do not provide fine discriminability in perceived depth. This means that the underlying tuning curves can be quite broad, reducing the number of units required to represent the full range of possible values along that dimension. Indeed, this may be the case for texture information: interpreted in light of the proposed model, the lack of change in the influence of texture information in Experiment 2 suggests that the tuning of network units may be broader along the texture dimension than along the stereo dimension. This broader tuning would inevitably cause increased interference among the units that encode the limited range of texture slants we tested [39], helping to explain why the influence of texture information did not change significantly in either condition. Second, the presumably large representational capacity of the visual system could be adaptively leveraged in a way that maximizes the number of units devoted to frequently observed combinations of depth cues, while minimizing the number that represent combinations that never occur. Even with these efficient encoding strategies, however, the representational capacity of the visual system would have to be extremely large to support the proposed combinatorial system.

### Implications of the double-pointing model of reach-to-grasp control

Turning away from models of cue reweighting, an important final discussion point involves our generic assumption that estimates of 3D shape properties (*i.e.,* surface slant and object depth) are, in fact, the inputs to the motor planning processes involved in our visuomotor tasks. In contrast to this assumption, Smeets and Brenner [40, 41] have notably defended an elegant alternative model of precision-grip control in which thumb and index finger movements are planned as two independent pointing movements aimed at two separate egocentric locations. Their model provides another possible way to explain the apparent stereo preference in visuomotor tasks involving 3D stimuli. According to their model, the visuomotor interactions in these studies are guided by egocentric distance estimates, which tend to rely strongly on vergence information from recent fixations (*i.e.*, stereo), and to be relatively insensitive to texture patterns and other pictorial information. In our experiments, however, we found that perceptually matched stimuli with different values of stereo slant and stereo depth elicited identical motor responses prior to sensory feedback (Figs 3, 5, and 8). Thus, we can conclude either (a) that our perceptual judgments were also based on egocentric distance estimates from multiple surface locations or (b) that 3D property estimates were in fact used for movement planning in our tasks. As a result, our main conclusions are unchanged under the double-pointing model of reach-to-grasp control.

## Conclusion

The operation of sensorimotor adaptation and cue reweighting over the short timescales tested in these experiments provides an alternative explanation of why some studies find a preferential reliance on stereo information in visuomotor tasks compared to perceptual tasks. Notably, this account avoids postulating separate encodings of 3D shape for perception and action. This broad theoretical conclusion is underscored by Experiment 3, where we found that cue reweighting involves correlated changes in perceptual judgments and motor responses. In contrast to the dissociated view of perception and action, these results suggest a close link between these two functions: distortions of 3D shape perception lead to improperly planned movements, but the resulting sensory feedback signals enable the system to compensate for those distortions.

## Methods

### Participants

Participants were between 18 and 35 years old, right-handed, and had normal or corrected-to-normal vision. They were either granted course credit or paid hourly as compensation. Informed consent was obtained from all participants prior to any participation. Our research protocol was approved by the Brown University Institutional Review Board and performed in accordance with the ethical standards set forth in the Declaration of Helsinki.

Fifteen participants were recruited for Experiment 1. Forty-eight participants were recruited for Experiment 2. Twenty-eight were exposed to the haptic-for-texture condition, and twenty were exposed to the haptic-for-stereo condition. One participant from the latter condition was excluded from analysis because more than half of their Grip Placement trials were marked for exclusion by the criteria indicated below. Fifty-six participants were recruited for Experiments 3A (N = 36) and 3B (N = 20).

### Apparatus

The general layout of the tabletop virtual reality apparatus used in Experiments 1 and 2 is depicted in Fig 2d; the layout for Experiment 3 is depicted in Fig 6a. Participants sat with the chin resting comfortably on a chinrest. Right-hand movements were tracked using an Optotrak Certus. Small, lightweight posts containing three infrared emitting diodes were attached to the index finger and thumb nails, and the system was calibrated to track the tip of each distal phalanx. This motion-capture system was coupled to a virtual reality environment consisting of an oblique half-silvered mirror that reflected the stereoscopic image on a 19” CRT monitor to provide consistent accommodative and vergence information at the intended viewing distance (very small, probably negligible, discrepancies between accommodation and vergence would arise when fixating points with non-zero disparity). The room was completely dark, and an opaque back panel was placed on the mirror to prevent vision of the hand or of the physical surfaces providing haptic feedback.

Participants viewed computer-generated 3D stimuli with stereo and texture cues controlled independently via backprojection. In Experiments 1 and 2, we rendered slanted surfaces that rotated around the transverse axis through the middle of the object, which was positioned at eye level at a distance of 38 cm. A frontoparallel surface (0° slant) diagonally subtended 13° of visual angle. In Experiment 3, object depth was based on a single cycle of a cosine function protruding toward the observer. The square bases of these “ridge” objects diagonally subtended 8° of visual angle viewed at a distance of 40 cm. In all experiments, the rendered 3D stimuli appeared to be floating in space beyond the mirror. Stereoscopic presentation was achieved with a frame interlacing technique in conjunction with liquid-crystal goggles synchronized to the frame rate. No visual feedback of the hand was provided in Experiments 1 or 2. In Experiment 3, small dots provided stereoscopic visual feedback of the index finger and thumb until one of the two fingers came within 25 mm of the target object.

In Experiments 1 and 2, haptic feedback was provided by a square plexiglass surface, attached to a stepper motor to control the slant and mounted on linear positioning stages to control the position. In Experiment 3, haptic feedback was provided by a custom-built motorized object consisting of a stepper motor with its shaft extended by a long screw, pointing toward the observer. Just beyond the tip of this screw, we mounted a plastic half-cylinder to simulate the rounded tip of the “ridge” stimuli. To simulate the rear side of the stimuli, a flat, wooden surface was threaded onto the screw, and two additional plastic half-cylinders were attached in positions corresponding to the top and bottom edges of the stimulus. As the stepper motor spun, this piece traveled back and forth along the length of the screw, anchored on one side so that one full rotation of the screw changed the haptic depth by one thread pitch. In all experiments, precise alignment between the physical objects and the rendered 3D stimuli was established at the start of each session. Before every trial, the state of the physical object was checked using additional Optotrak markers and corrected if necessary.

### Procedure

#### Experiment 1

Experiment 1 involved two tasks. First, in the Matching task, participants matched through a psychometric procedure the perceived slants of three stereo-texture conflict surfaces (*s_T_i__*, *s_S_i__*) with cue-consistent surfaces (smatch_i_, smatch_i_) where stereo and texture specified the same slant, such that *Ψ*(*s_T_i__*, *s_S_i__*) = *Ψ*(smatch_i_, smatch_i_). The three stimuli were chosen such that, for each surface, stereo slant and texture slant differed by a constant conflict angle of 30° (*s_T_i__* = *s_S_i__* + 30°). These combinations of stereo and texture slant are illustrated in Fig 2b. On each trial, we allowed participants to switch freely between the fixed cue-conflict stimulus and an adjustable cue-consistent stimulus, using keypresses to make incremental changes to the slant of the cue-consistent stimulus until it appeared to match the slant of the cue-conflict stimulus. To prevent the use of motion information, we displayed a blank screen with a small fixation dot for an inter-stimulus interval of 750 ms whenever the stimulus was changed. Participants performed five repetitions of matching for each of the three fixed cue-conflicts, for a total of 15 trials.

The resulting set of six stimuli (three pairs of matched cue-conflict and cue-consistent surfaces) were then presented as stimuli in the Grip Placement task. With the hand shaped into a precision grip, participants reached toward the displayed surface with the goal of making the index finger and thumb hit the surface at the same time. They were instructed to use only the wrist and arm to change the grip orientation while holding their fingers in fixed precision grip posture. This task is similar to handheld object placement tasks where cylinders are placed so that their bottoms are parallel with the surface at contact (Knill, 2005), but also engages the fingers’ tactile sensitivity. A standard, three-phase “ABA” design was adopted for the visuomotor task. In the Baseline phase, participants reached toward their personalized set of cue-consistent surfaces for 30 trials. At the transition from Baseline to the Adaptation phase, the cue-consistent surfaces were suddenly replaced by the perceptually matched cue-conflict surfaces, with the underlying physical surface reinforcing the texture slant. Following 180 trials of exposure to these conflict surfaces, the experiment concluded with a 30-trial Washout phase, identical to Baseline. Throughout this task, we used a binned trial order such that each of the three surfaces was presented once before any one was repeated, facilitating local averaging of trials.

#### Experiment 2

As in Experiment 1, participants in Experiment 2 again performed the Matching and Grip Placement tasks, but there were nine target stimuli presented during Adaptation (Fig 4a): three cue-consistent (main diagonal) and six cue-conflict (off-diagonal cells). The six cue-conflict surfaces were perceptually matched with adjustable cue-consistent stimuli in the first block using the same psychometric procedure as Experiment 1. We eliminated the Washout phase from this experiment in order to reduce the overall duration. During Baseline, the targets were the nine cue-consistent perceptual matches, specific to each subject. These nine targets were presented four times each in a binned trial order for a total of 36 Baseline trials. During Adaptation, the visual stimuli were the nine target stimuli, and the physical surface slants were consistent either with the texture slants (haptic-for-texture group) or the stereo slants (haptic-for-stereo group). These were presented 14 times each in a binned trial order for a total of 126 trials.

#### Experiment 3

The procedure of Experiment 3 is generally similar to Experiments 1 and 2, but we made a number of adjustments in order to facilitate direct comparison of perceptual and motor task data. In particular, the relevant stimulus set involved depth metamers, which were indicated by each participant in a Pre-test, and again in a Post-test. During each Pre-test and Post-test trial, participants were repeatedly shown (for 750 ms each) the 30-mm cue-consistent standard, followed by a random dot cloud filling a 5-cm cubic volume (a mask that also served to prevent fatigue effects), followed by the adjustable cue-conflict stimulus with their current stereo depth setting. The adjustable cue-conflict had one of four fixed texture depths (18, 24, 36, or 42 mm). Participants used keypresses to incrementally change the stereo depth of the cue-conflict object until they indicated a perceptual match. Every keypress resulted in the two objects being displayed again in sequence as described above. Participants performed six repetitions with each of the four cue-conflicts, for a total of 24 trials.

The second key methodological difference with Experiments 1 and 2 is that we changed from the Grip Placement task to a Grasping task, where participants applied a front-to-back precision grip to the targets. With the fingers pinched closed at the start position, participants viewed the object for 500 ms plus a random jitter of 0-100 ms prior to receiving a “go” signal. Participants then had 2 seconds to successfully complete the trial. To successfully complete a trial, participants were required to place the thumb on the plastic contact at the front tip of the object, the index finger on the plastic contact at the top-rear edge, and to hold still for 300 ms. If they did so within the allotted time, a pleasant feedback tone was played, otherwise an aversive buzzing noise was played.

In the Adaptation phase of Experiment 3A, we presented the personalized depth metamers from the Pre-test, which were paired with haptic depths of 18, 24, 30, 36, and 42 mm. In the haptic-for-texture condition, haptic depths matched the texture depths, and were therefore inversely correlated with the stereo depths. In the haptic-for-stereo condition, the five haptic depths were matched up in the opposite order across the metamer set; thus, haptic depths were positively correlated with stereo depths, but did not exactly match them, and were negatively correlated with texture depths. To provide some variation in perceived depth, we replicated the primary set of five metamers twice more by shifting the stereo and texture depths for each metamer by 2D vector distances of ±4 mm, perpendicular to the best-fit line through their Pre-test data; the corresponding haptic feedback was also shifted by ±4 mm. These fifteen objects were presented in seven Adaptation bins for a total of 105 Adaptation trials. In this full set of target objects (pooling over participants), the Pearson correlation coefficients (*r*) between haptic depth and stereo depth were +0.87 and −0.31 in the haptic-for-stereo and haptic-for-texture conditions, respectively. The correlation coefficients between haptic and texture depth were −0.76 and +0.93, again respective to the two conditions.

Prior to the Adaptation phase of Experiment 3A, participants grasped a slightly different set of objects in a 26-trial Baseline phase. In the first 13 trials, we presented cue-consistent stimuli with depths of 18, 24, 30, 36, and 42 mm. The 30-mm stimulus was repeated five times while the others were repeated twice. In the latter 13 trials, we retained the 18, 24, 36, and 42-mm cue-consistent stimuli, again presenting them two times each, but the five 30-mm cue-consistent stimuli were replaced by the participant’s personalized set of five depth metamers. During these first five presentations of the metamers, the underlying haptic depth was always 30 mm.

In the Adaptation phase of Experiment 3B, we repeatedly presented five cue-conflict objects with a constant 8-mm difference between texture depths (18, 24, 30, 36, and 42 mm) and stereo depths (10, 16, 22, 28, and 36 mm). The associated haptic depth always matched the texture depth. Participants grasped these five objects over 21 bins for a total of 105 Adaptation trials. Prior to the Adaptation phase of Experiment 3B, we included a 27-trial Baseline phase, composed of three bins of nine grasps toward cue-consistent stimuli spanning from 18 to 42 mm in 3-mm increments.

### Analysis

Raw motion-capture position data was processed and analyzed offline using custom software. Missing frames due to marker dropout were linearly interpolated, and the 85-Hz raw data was smoothed with a 20-Hz low-pass filter. We excluded from analysis all trials where (a) the proportion of missing frames exceeded 90%, (b) fewer than 5 frames were not missing, or (c) the center of the grip traveled less than 25 mm. In Experiment 1, these criteria resulted in the exclusion of 79 individual trials out of a total 3,600 trials. In Experiment 2, these criteria resulted in the exclusion of 298 out of a total 7,776 trials. In Experiment 3A, three out of a total 9432 trials were excluded by these criteria. In Experiment 3B, five out of a total 2640 trials were excluded by these criteria.

In Experiments 1 and 2, in-flight grip orientation was calculated as the declination of the projection of the line joining the fingertips onto the sagittal plane. To infer the planned grip orientation at the end of the movement, we used a snapshot of the grip orientation at a moment 10 mm prior to contact (Fig 11a). This was done in order to avoid any contamination caused by adjustments made after one of the fingers contacted the surface. To extract this *terminal* grip orientation, we first processed the entire trajectory, then searched for the first motion-capture frame where either one of the fingers contacted the physical surface (minimum orthogonal distance from each finger to the surface plane), then scanned 10 mm backward along the trajectory.

**Fig 11.**
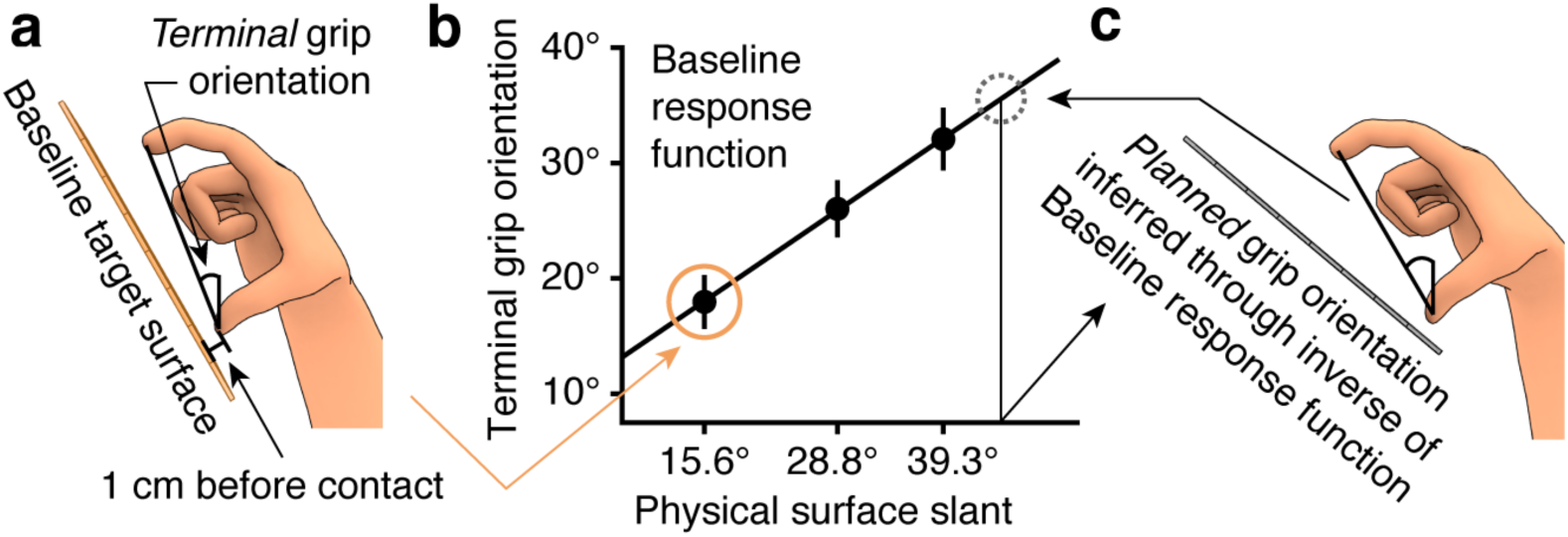
Method for estimating planned grip orientation based on terminal grip orientations. See text for details.

However, this terminal grip orientation does not represent the planned grip orientation because the movement is still in progress. For the data displayed in Fig 3c (Experiment 1), we used each participant’s Baseline performance to determine how their measured terminal grip orientations related to the actual physical slants they intended to place the fingers on, which were presumably their planned grip orientations (Fig 11b). Accordingly, we fit a linear regression to this Baseline data and used the inverse of the estimated function to transform terminal grip orientations, our raw dependent variable, into the planned grip orientations (Fig 11c). This does not affect the statistical analysis or the modeling of our data, but simply helps to present the data in a more understandable format, with the dependent variable sharing the same metric as the rendered slant information.

In Experiment 1, we fit the error-correction parameter *b* of our first adaptation model to minimize the root mean squared error of the model with respect to the average trial-by-trial grip orientations. To do so, we used the constrained optimization by linear approximation (COBYLA) algorithm (Powell, 1998) of the nloptr package (Johnson, *n.d.*) in R (R Core Team, 2014), constraining the fit so that *b* was bound between 0 and 1. After fitting the model, we observed that participants’ planned grip orientations converged on a slightly greater value than did the model, as the model was bound to converge on the actual physical slants that were presented. In order to incorporate this empirical measure of the fully adapted state into the model before making predictions for the Washout phase, we manually set the internal state of the model on the first trial of Washout to reflect the average change in grip orientations from Baseline to the final 30 trials of Adaptation.

The factorial design of the conflict stimuli in Experiment 2 allowed us to measure the relative influence of stereo and texture information in the Grip Placement task by estimating coefficients (slopes) for each cue via multiple linear regression according to Equation 5, with the terminal grip orientation as the response variable. Unlike in Experiment 1, we did not transform the terminal grip orientations into planned grip orientations—this was not necessary in Experiment 2 because we were interested only in how the slope coefficients from the multiple regression changed over time. Just as before, terminal grip orientations were measured by finding the moment at which one of the fingers first touches the surfaces, then scanning backwards by 10 mm of hand movement. A regression was computed for each bin of nine trials within the Adaptation phase, producing a fine-grained timeline of the influence of each cue on terminal grip orientation. In Baseline, we computed the slope of the terminal grip orientation with respect to the perceptually matched slant values using simple linear regression in each bin.

In Experiment 2, since there were no differences in procedure between the two feedback conditions until the Adaptation phase of the Grip Placement task, and since no significant differences were found in Matching performance between the two groups for any of the six cue-conflict stimuli, the two groups were combined when analyzing the Matching task.

In Experiment 3, the relative cue weights in the Pre-test and Post-test were computed based on the stereo and texture settings for the four cue-conflict metamers. Since the cue-consistent standard stimulus had a depth of 30 mm, the relative influence of stereo information can be computed as *w_S_* = (30 − *z_T_*)/(*z_S_* – *z_T_*), where *z_T_* and *z_S_* denote, respectively, the fixed texture depth and the final stereo setting. This equation implies a weighted linear combination of the two cues where the weights sum to one, so the texture weight is obtained by subtracting the stereo weight from one.

To analyze the grasping data of Experiment 3A, we fit linear regressions to the MGAs in each bin as a function of the texture depths across the metamer set. To analyze the correlation between perceptual and motor changes, we transformed the change in relative cue weights for each participant into a prediction of the MGA slope across the five metamers that corresponds to the observed perceptual change: *k_metamers_* = ((1 − *w_S_Post__*) – (1 − *w_S_Pre__*)) / *w_S_Pre__*), where *w_S_Pre__* and *w_S_Post__* are the stereo weights in Pre-test and Post-test, and *k_metamers_* is the predicted MGA slope across the metamers. Empirical MGA slopes were computed by fitting a regression to the average MGAs from the entire Adaptation phase as a function of the texture depths across the metamer set. In Experiment 3B, we predicted the decrease in the average MGA at the beginning of the Adaptation phase by multiplying the Pre-test stereo weight by the imposed 8-mm decrease in stereo depth, then subtracting this value from the average MGA in the final Baseline bin.

## Acknowledgments

We thank Carlo Campagnoli for numerous helpful discussions and assistance with figure preparation.

